# Novel correlative microscopy approach for nano-bio interface studies of nanoparticle-induced lung epithelial cell damage

**DOI:** 10.1101/2024.10.11.617573

**Authors:** Rok Podlipec, Luka Pirker, Ana Krišelj, Gregor Hlawacek, Alessandra Gianoncelli, Primož Pelicon

## Abstract

Correlated light and electron microscopy (CLEM) has become essential in life sciences due to advancements in imaging resolution, sensitivity, and sample preservation. In nanotoxicology— specifically, studying the health effects of particulate matter exposure—CLEM can enable molecular-level structural as well as functional analysis of nanoparticle interactions with lung tissue, key for the understanding of modes of action. In our study, we for the first time implement an integrated high-resolution fluorescence lifetime imaging microscopy (FLIM) and hyperspectral fluorescence imaging (fHSI), scanning electron microscopy (SEM), ultra-high resolution helium ion microscopy (HIM) and synchrotron micro X-ray fluorescence (SR µXRF), to characterize the nano-bio interface and to better elucidate the modes of action of lung epithelial cells response to known inflammatory titanium dioxide nanotubes (TiO₂ NTs). Morpho-functional assessment uncovered several mechanisms associated with the extensive DNA, essential minerals and iron accumulation, cellular surface immobilization, and the localized formation of fibrous structures, all confirming immunomodulatory responses. These findings advance our understanding of the early cellular processes leading to inflammation development after lung epithelium exposure to these, high-aspect-ratio nanoparticles. The novel experimental approach, exploiting light, ion and electron sources, provides a robust framework for future research into nanoparticle toxicity and its impact on human health.

## Introduction

Among the novel microscopy approaches, correlated light and electron microscopy (CLEM) has attracted considerable interest in the life sciences community, particularly following recent developments in the spatial resolution and sensitivity capabilities of individual techniques.^1,2^ Advanced live cell imaging, with its targeted labelling of individual cellular components, can provide molecular specificity and the observation of dynamic changes.^3,4^ In addition, the advanced imaging of fixed cells, when performed in high vacuum, can provide ultrastructural information with nanometer spatial resolution ^5^ and chemical information with sub-micron resolution.^6,7^ When successfully combined on the same region of interest (ROI), a correlative microscopy approach offers several significant advantages. It provides comprehensive functional and structural insights at multiple scales, as well as increased accuracy through cross-validation of data and improved contextual understanding through the analysis of dynamic and static processes on live and fixed cells, respectively.

One of the highly relevant scientific areas that can be addressed by correlative microscopy is the field of nanotoxicology, which studies the environmental and, in particular, health effects of the ubiquitous particulate matter in the polluted air.^8^ In particular, ambient ultrafine particles (UFPs) and engineered nanomaterials (NMs) less than 100 nm in size can penetrate deep into the lungs and induce a variety of chronic diseases, including inflammation, fibrosis and cancer. ^9–11^ For safety assessment and prediction of nanoparticle-induced adverse outcomes, understanding of the mode of action at the molecular level, starting from the moment the particles interact with the lung tissue, is critical. To this end, several innovative correlative microscopy approaches have been introduced in recent years to elucidate the interaction at the nano-bio interface and the subsequent impact of nanomaterials and/or elements on biological matter.^12–36^ A commonly used approach is CLEM^12^ which mostly combines highly specific and targeted fluorescence microscopy (FM) with transmission electron microscopy (TEM).^13–15^ More recently, super-resolution fluorescence imaging combined with TEM and scanning electron microscopy (SEM) has been implemented to address the complex biological questions at the nanoscale.^16^ While CLEM remains a prominent approach, recent years have seen the development of several complementary techniques. These correlate FM with X-ray tomography^17^, dark-field microscopy (DFM)^18^, atomic force microscopy (AFM)^19,20^, and ion-induced luminescence.^21^ Additionally, to enhance and support CLEM, a few ion-based methods have also been introduced.^22,23^ For more detailed and quantitative analysis of the specific molecules and elements present in biological tissues, correlative microscopy approaches often include secondary ion mass spectrometry (SIMS) combined with electron microscopy^24–26^ and novel high-resolution helium ion microscopy (HIM).^27–29^ In addition to SIMS, proton-induced X-ray emission (PIXE) and X-ray fluorescence (XRF) analytical techniques are also used to quantify chemical elements in biological samples, and are combined with FM^30^ and EM^31,32^, respectively. Other correlative approaches include the combination of XRF with AFM^33^ or XRF with ion beam techniques^34–36^ for elemental quantification and complementary information.

Despite the continuous development of new correlative microscopy approaches in nanomaterial toxicity research, the full potential of multimodalities and their combinations has not yet been fully exploited, with many opportunities and challenges.^37^ In this study, we present a novel correlative microscopy pipeline that can investigate the modes of action and toxic effects of UFPs and NMs, both of which pose the greatest risk to the lungs, as they can penetrate deep into the alveolar region and potentially further into the bloodstream.^38^ We are particularly focused on the worldwide and one of the most widely used industrial NMs, titanium dioxide (TiO_2_), which in the form of high surface area nanotubes (NTs) induces lung inflammatory^39,40^ and is likely to be carcinogenic for humans.^41^ We demonstrate the utility of an innovative multimodal and multiscale experimental workflow that integrates several complementary imaging and spectroscopy techniques on live and fixed cells in a high vacuum (Figure 1). In order to preserve the nanoscale morphology and properties of the investigated nano-bio interfaces on the surfaces, we use a rapid cryofixation technique without chemical cross-linking, which could otherwise compromise sample size, shape or cell membranes at the investigated spatial scale.^42–46^ We must acknowledge that the development of sample preparation and preservation for high-vacuum, high-resolution correlative microscopy is one of the major challenges in the field and can play a key role in influencing the physicochemical properties of samples and the subsequent interpretation of results.^7,33,34,37,47^

**Figure 1.**
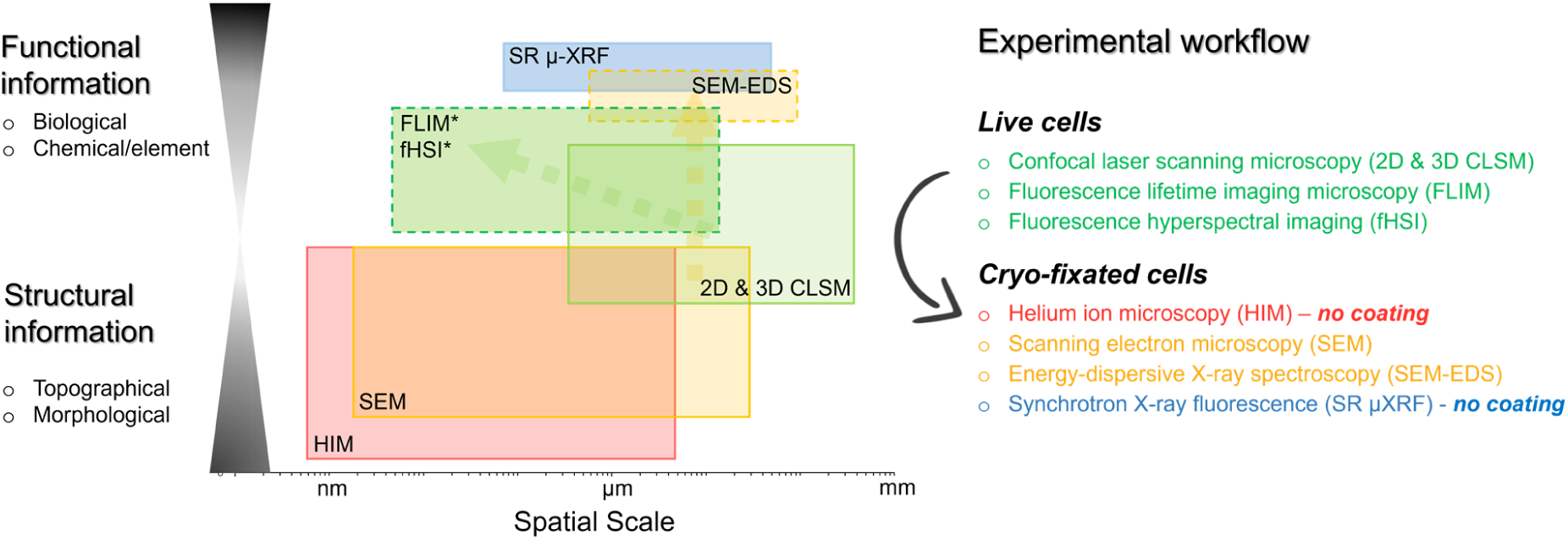
Novel multimodal microspectroscopy-based correlative microscopy enables a comprehensive investigation of the studied nano-bio interface across a broad spatial scale, from tens of microns down to the nanometer level. This approach provides both functional and structural insights, which in our study focus on the interface between lung epithelial cells and exposed NMs. The colored areas illustrate the typical imaging ranges for each technique, highlighting the type and extent of information obtained. * The spatial resolution extends beyond the optical limit, as FLIM and fHSI can probe molecular interactions at a sub-diffraction scale.

The present study aims to cover a wide range of detectable spatial scales (down to the nanometer level) and to gain insight into both functional and structural properties of the samples under investigation. In order to obtain comprehensive knowledge, data gathered from multimodal fluorescence microspectroscopy performed on living cells were integrated with data obtained from high resolution and high elemental sensitivity X-ray, ion and electron-based techniques performed in high vacuum. These approaches provided new insights into the physicochemical properties of cell-excreted and cell-deposited TiO_2_-biological matter (TiO_2_-bio) composites aggregated and immobilized on the surface of the lung epithelium. In addition, we gained deeper morpho-functional insights into the inflammatory and anti-inflammatory mechanisms triggered by the binding and transport of DNA and other molecules to TiO₂ NTs. We also uncovered the possible mechanism of an initiation of an acute cellular response via the formation of a fibrous network over the nanoparticles. These findings contribute to a deeper understanding of the current knowledge in the field of UFPs and NMs-induced toxicity on the lung epithelium.

## Results and discussion

### Impact of TiO_2_ NTs on DNA binding and the transport in apoptotic lung epithelial cells

To gain new insights into the response of lung epithelial cells to exposed metal oxide NMs, we first performed an integrated multimodal high-resolution fluorescence imaging on live cells exposed to TiO₂ NTs for two days with a meaningful surface dose (3 µg/cm^2^), as discussed in Supplementary Comment #1 (Figure 2A-B). Cells were grown on special, 10 nm thin Finder grid Formvar/Carbon substrates, with fiducials used for correlative microscopy (Figure 2A, arrow). Prior to measurement, the observed cell mitochondria, actin network, and nuclei were stained to characterize any potential structural and/or functional changes that may have occurred as a result of nanoparticle exposure. To track and determine potential co-localization between stained cellular structures and nanoparticles, TiO₂ NTs were fluorescently labeled prior to administration according to the well-established protocol.^48^ To be able to successfully register and overlap the correlated images by identifying the fiducials, we first applied low-magnification imaging (10X) using two-channel confocal laser scanning microscopy (CLSM). In addition to clearly visualizing the grid, used as a fiducial (Figure 1A, in black), it was possible to distinguish between TiO₂ (in red) and mitochondria (in green). To delve deeper into the interaction between TiO₂ and cells, we performed high-resolution imaging in three dimensions (3D) at 60X magnification (Figure 1A, second image). The 3D image shows the localization and aggregation of TiO₂ on the cell surface (white arrow), a phenomenon observed and described in our recent study.^39^ Here, we further unravel the high heterogeneity of the composites in morphology and size (upper image plane (UP)). However, the similar excitation and emission profiles of the fluorescent dyes prevented a clear differentiation between the TiO_2_, nucleus and actin structures beneath the cell membrane (lower image plane (LOW)), using a 2-channel CLSM.

**Figure 2.**
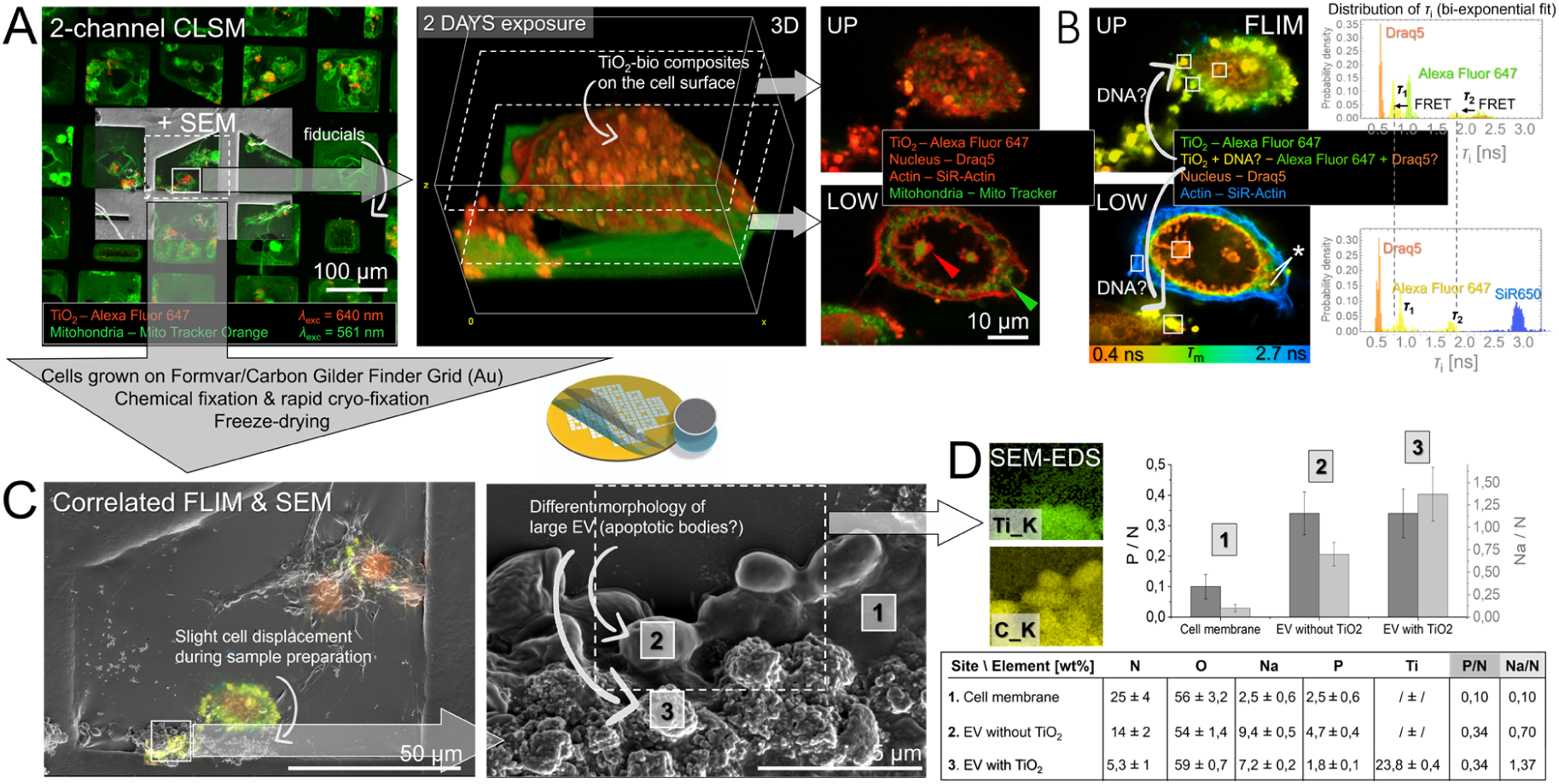
Correlative multimodal fluorescence microscopy and SEM-EDS (multimodal CLEM) applied on lung epithelial LA4 cells exposed to TiO_2_ NTs for 2 days reveals physico-chemical properties of the nano-bio interface and the insights of DNA binding and transport during plausible cell apoptosis. A) Low to high magnification 2D/3D CLSM showing TiO_2_-bio composites on the cell surface and the morphology of the cell nuclei, actin and TiO_2_ (in red) and mitochondria (in green). B) The same structures with correlated FLIM capable of precisely distinguishing between individual labeled structures (color coded from red to blue),) using single laser source (*λ* = 640 nm). Shifted distribution of *τ*_i_ and *τ*_2_ from bi-exponential fit (green to yellow in the chart on the right) indicate on local FRET between TiO_2_ and DNA dyes, revealing plausible binding of DNA to TiO_2_ surface. C) Correlated FLIM and SEM after a successful registration and overlap of fiducials shown in A). High magnification image reveals large EVs with different morphologies, denoted with 2 and 3. D) Chemical characterization of the same structures performed by SEM-EDS confirms not only high accumulation of biological matter (C) on nanoparticles’ surface (Ti), but also a few times higher content of organic phosphate normalized to biological matter (P/N) (right chart and table) indicating a high but mechanistically different DNA load inside large EVs.

To identify these individual structures, we proceeded with fluorescence lifetime imaging microscopy (FLIM), integrated into the same experimental setup. FLIM, a powerful imaging technique for measuring the lifetime of fluorescent molecules independent of signal intensity,^3^ also provides quantitative insights into the properties and dynamic changes of the surrounding molecular environment^49^ - an advantage we later utilized. With FLIM, we were able to identify the individual labeled structures (Figure 2B), which was not possible with the 2-channel CLSM. The fluorescence lifetime, as measured in each image pixel, is color coded with the mean lifetime (*τ*_m_), which is the weighted average of the individual lifetime components (*τ*_i_) from the bi-exponential fit (see Equation 1). The FLIM analysis, performed after using a single excitation laser (*λ* = 640 nm), allowed to distinguish between actin structures (shown in blue), a nucleus undergoing a distinct phase separation due to specific biophysical and/or biochemical processes^50^ (shown in orange), and TiO₂ with a wide *τ*_m_ distribution (shown in green to yellow). The latter indicates on a high degree of structural and molecular variability and most likely local attachment of DNA molecules and/or their fragments to the surface of nanoparticle composites (see the regions indicated by the arrow). The proximity of the Draq5 dye, intercalated in the DNA structure at a distance of less than 10 nm, to the Alexa Fluor 647 dye attached to TiO_2_ plausibly induces a fluorescence resonance energy transfer (FRET) transition, which is known to shorten the fluorescence decay time.^51^ This is evidenced by the blue shift in both the *τ*_1_ and *τ*_2_ distributions (Figure 2B, upper right image). We also observed the Draq5 signal locally concentrated in the cell cytoplasm (see the asterisk). With the additional experiments taking advantage of the combined FLIM and fluorescence hyperspectral imaging (fHSI), we further confirmed the local and widespread presence of DNA in the cytoplasm, apparently co-localized with the TiO_2_ NTs (Figure S1), suggesting a possible downstream inflammation.^52^ These intracellularly formed nano-bio composites may represent one of the key early states in the cell clearance mechanism that evolves into the observed surface-immobilized TiO_2_-bio composites.

To gain further insight into the structural and functional properties of the formed composites in relation to DNA binding, we then prepared the samples for correlative SEM (Figure 2C). Due to the chemical fixation treatment of 4% PFA and 2% GA and the fragile thin formvar/carbon support films, we observed a slight displacement of the investigated cell and its structures as indicated by the white arrow. However, this was not a limitation in this study, as the identification of the same extracellular vesicles (EVs) in fixed cells was sufficient for subsequent morphological and chemical characterization (Figure 2C-D). The SEM measurements revealed the presence of distinct morphological features of large EVs that resembled apoptotic bodies (ABs).^53^ Additional SEM-EDS chemical mapping (Figure 2D) with typical spectra (Figure S2) revealed that rough surface topography was co-localized with the presence of TiO_2_ (Ti map in green), while all large EVs exhibited a similar amount of organic matter (C map in yellow). Given that the electron penetration depth and interaction volume for the detected characteristic X-rays are on the order of a few microns at E₀ = 15 keV,^54^ and considering the much thinner measured layer of organic matter in EVs with a high nanoparticle content (Ti 23.8 wt%), we confirmed a concentrated accumulation and extensive binding of organic matter (lipids and proteins) to the TiO₂ surface— consistent with findings from our recent study.^55^ Through detailed chemical analysis of the distinct regions labeled 1-3 (Figure 2C), we confirmed previous FLIM-based observations suggesting the very likely presence of DNA in EVs (Figure 2D). The weight percentage (wt%) of the elements shown in the table and in the corresponding bar charts show a clear, more than threefold, elevation of phosphate (P) and sodium (Na) inside EVs (sites 2 and 3) compared to the surrounding cell membrane (site 1). The values are normalized to the concentration of organic matter - in our case organic nitrogen (P/N and Na/N) – within each measured region to remove the potential effects of different densities and interaction volumes, all of which affect EDS detection. It is noteworthy that there was no discernible difference in P/N between EVs with high and negligible TiO_2_ content, indicating a similar concentration of DNA or its fragments in both cell-excreted structures. However, the different morphologies of these large EVs, observed by both high magnification SEM and FM on live cells, suggest on completely different functions and modes of action for DNA loading, packaging and transport, all of which are considered essential for cellular homeostasis and immune response modulation.^56^

The most likely cause of the high P accumulation in flat-surfaced EVs (site 2) is the high concentration of DNA with associated binding proteins in the so-called plasma membrane blebbing process, followed by its release from the cells in the form of apoptotic bodies, being the largest class of EVs. This has the biological implication of impending apoptosis of the measured cell, with the shape indicating still on an early stage of this process. However, the initiation of cell apoptosis may well be indicated by the observed chromatin condensation, which is known to be one of the sequences of nuclear changes culminating in cell apoptosis,^57^ and the observed round mitochondria, which are known to occur under conditions that compromise mitochondrial function, such as cell apoptosis^58^ (Figure 2A, see arrows in lower right image). On the other hand, the EVs with a rough surface and a significant accumulation of TiO₂ appear significantly different from the previously observed EVs, casting doubt on whether these structures are truly apoptotic bodies. The most likely cause of the high concentration of organic phosphates (P), further supported by the highest accumulation of mineral Na^+^ among the measured sites, is a physical rather than a biological mechanism, as previously reported.^59,60^ Namely, the DNA phosphate backbone has been found to play the major role in TiO_2_ adsorption ^59^, while Na^+^ has been found to preferentially adsorb to the TiO_2_ surface as inner-sphere complexes,^60^ both under high ionic strength conditions, ^59^ such as our cellular environment. Preferential DNA binding to TiO_2_ surface, being found in both cytoplasmic (Figure S1) and cell surface region (Figure 2), may have important biological implications for disturbing cellular homeostasis. Interference with transcription or even replication processes, as observed in our recent study,^41^ may lead to genotoxic effects.^61^ On the other hand, Na^+^ binding, if carried out intracellularly, could disrupt the ion homeostasis, leading to disruption of membrane potential and the related processes. Moreover, the immobilization of TiO_2_-rich, irregularly shaped EVs, on the cell surface by the intertwined actin fibers from the inside (Figure S3, super-resolution STED microscopy on live cells) and fibrin-like fibers from the outside reported in Figure 5, could affect the normal clearance of apoptotic cells and potentially introduce pathological conditions for the inflammation or autoimmunity.

To elaborate on this hypothesis, the observed, dense structure of immobilized EVs on the apoptotic cell surface could partially block the secretion pathways of the typical anti-inflammatory signals, thereby disrupting the typical anti-inflammatory environment. Moreover, if apoptotic cells are not cleared efficiently and in a timely manner by macrophages due to their overload in high particulate matter exposure and plausibly reduced motility,^62^ they may progress to secondary necrosis and lose their membrane integrity.^63^ By releasing immunostimulatory content or so-called damage-associated molecular patterns (DAMPs), the cellular dynamics then promotes inflammation and recruitment of immune cells. DAMPs are released through the lesions in the plasma membrane orchestrated by the plasma membrane protein Ninjurin-1 (NINJ1), which is structured into supramolecular filamentous assemblies as recently shown by super-resolution microscopy.^64^ A meaningful observation is that the polymerized filaments range in size from a few tens to a few hundred nanometers—closely matching the dimensions of the lesions or holes we found on the cell surface close to the anchored TiO₂-rich excreted structures (Figure 3D). Thus, the holes may not necessarily be entirely a consequence of the incomplete sample preservation during cryofixation and freeze-drying, but may actually represent the plasma membrane lesions. However, further studies with better statistics are needed to confirm this.

**Figure 3.**
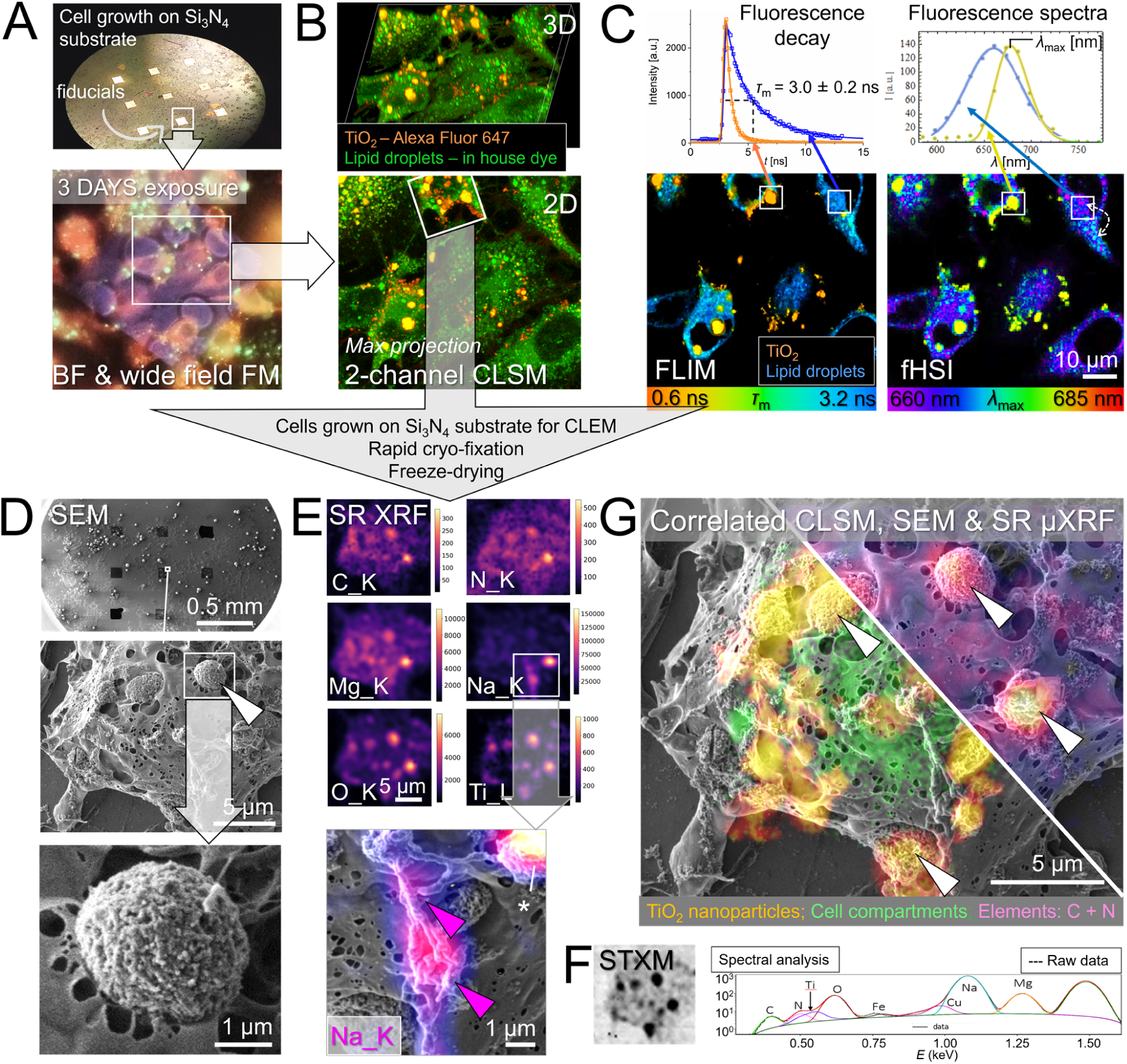
Novel multimodal CLEM combined with synchrotron µXRF (CLEM-µXRF) revealing in-depth physico-chemical and morpho-functional properties of inflammatory TiO_2_ NPs-rich cell-excreted composites immobilized on the surface of lung epithelial LA4 cells. A) Combined BF and wide field FM employed to visualize cell growth on 100 nm thick transparent silicon nitride (Si_3_N_4_) films of the 100*100 um dimension used as fiducials and optimal substrate for the correlative microscopy both reflection and transmission mode (µXRF). B) 2D and 3D CLSM performed on the same site to show labeled cells (in green) and cell-excreted TiO_2_ composites (in orange) with an improved resolution. C) The same site characterized with additional FLIM and fHSI, capable of distinguishing between individual labeled structures and of inspecting local microenvironment differences, such as molecular conformations or polarity. The upper charts illustrate FLIM decay curves and fluorescent spectra for the two distinct sites. D) Low to high magnification SEM images of the same site performed after rapid cryo-fixation of the sample to preserve distinct structural features of the TiO_2_-bio composites on the cell surface (bottom image). E) SR µXRF elemental maps from the same site showing the extensive binding of intrinsic biomolecules (C and N) and essential minerals (Na^+^ and Mg^2+^) on the TiO_2_-rich local sites. High-magnification correlated Na^+^ map and SEM further highlighting very likely functional relationship between composition and filamentous structure with potential inflammatory implications. F) Supporting atomic number (Z) and density-sensitive STXM map with the typical XRF spectra collected from the TiO_2_-bio composite site (denoted with an asterisk) and its color-coded fits for individual elements. G) Correlated live cell CLSM and high-vacuum SEM and µXRF revealing morpho-functional properties on the distinct sites (white arrows).

Taken together, this provides an additional possible mode of action in the initiation of chronic lung injury—complementing the well-established mechanisms of PM- and TiO₂-induced macrophage phagocytosis dysfunction^65–67^ and our recently discovered nanomaterial cycling between the lung alveolar surface and macrophagesl,^39^ both of which lead to chronic inflammation.

### In-depth morpho-functional characterization of cell-excreted and surface-immobilized TiO_2_-bio composites

To further investigate the morphological and structural properties of TiO₂-bio composites immobilized on the cell surface, as well as to gain further insight into their chemical properties at the sub-micron scale, we implemented an extended approach, a multimodal CLEM combined with synchrotron µ-XRF (CLEM-µXRF), where we joined 2-channel CLSM, FLIM and fHSI on living cells with SEM and µ-XRF on cryo-fixated cells (Figure 3). To improve the quality of the imaging capability and sample preservation in our novel correlative microscopy pipeline, we used special 100 nm thick silicon nitride (Si_3_N_4_) substrates/windows (Figure 3A) that are transparent to soft X-rays and avoided chemical fixation to preserve the nanoscale features of the investigated nano-bio structures (Figure 3D), respectively. In order to improve the resolution and sensitivity of the analysis of particularly light elements, we integrated spatially resolved synchrotron radiation SR µXRF (Figure 3E-F), with its ability to achieve up to an order of magnitude better, sub-micron spatial resolution and up to several orders of magnitude better elemental sensitivity^68^ than SEM- EDS, making it a valuable tool for investigating intricate biological processes such as ours.

Lung epithelial cells were grown on a surface modified Si_3_N_4_ substrate with fiducials in the form of multiple 100 x 100 µm transparent windows and exposed to fluorescently labelled TiO_2_ NPs for a period of 3 days. We observed a large number of TiO_2_-rich structures, measuring up to a few microns in size as shown in green and orange using a wide-field FM (Figure 3A) and enhanced-resolution CLSM (Figure 3B), respectively. For visualization, the cells were stained with the in-house synthesized lipid droplets (LDs) dye. Due to partitioning of the dye to other lipid-rich environments during the incubation period, the contrast of the LDs was reduced, as shown by the green color (Figure 3B). With CLSM, in comparison to a wide field FM, we were able to better distinguish each of the numerous TiO₂ NTs-rich composites, which varied in size and shape.

To gain further insight into the local properties and interactions in the near molecular environment of the observed TiO₂ NTs-rich structures and LDs, we performed FLIM and fHSI imaging/mapping at the same site (Figure 3C). The typical and easily distinguishable FLIM and fHSI spectra from the two distinct cell compartments within the white rectangles and corresponding fits using a bi-exponential function (Equation 1) and an empirical intensity-normalized log-normal function with nanometer peak position resolution,^69^ respectively, are shown in the top. The fHSI of LDs and lipid-rich regions showed spectral shifts of up to 10 nm, ranging from *λ*_max_ = 660 nm (in purple) to *λ*_max_ = 670 nm (in blue), indicating local microenvironmental differences, such as molecular conformation, polarity, etc.^70^ This was supported by FLIM data, with LDs showing a reduced mean fluorescence lifetime (*τ*_m_) (lighter blue) compared to other lipid-rich regions (darker blue), indicating an increased packing density of the dye within LDs. The analysis of TiO₂ NPs-rich structures showed less heterogeneity, but still with a considerable range in *τ*_m_, ranging from 0.6 to 1.0 ns, also indicating on differences in the packing density. An increased *τ*_m_ may also indicate a greater distance between fluorophores, possibly due to lipid interspacing, as shown previously.^39^

To probe deeper into the properties and possible consequent effects of the TiO₂ NTs-bio interaction, we introduced to our pipeline an additional correlated SEM and SR µXRF (Figure 3D- E). The combination of these advanced techniques provided new insights into the morphological and structural features well below the micron scale. The use of a 150 mM ammonium acetate wash solution to remove buffer salt crystals, which otherwise interfere with structural and chemical analysis,^71^ followed by rapid cryofixation by plunge freezing and final lyophilization, allowed the preservation of the majority of cell surface structures, including the studied TiO_2_ NPs-rich excreted composites studied (Figure 3D). The high magnification SEM image shows an almost perfect spherical structure (indicated by the white arrow), likely resulting from the most efficient packing of nanoparticles with the lowest surface energy-maximizing stability.^72^ To confirm that the structure is composed of nanoparticles, we observed strong co-localization with the fluorescence signal of a TiO₂ dye, as measured by CLSM in live cells (Figure 3B, outlined in a rectangle). This is further illustrated by the overlap of all correlated results from CLEM-µXRF in Figure 3G (indicated by the arrows). To obtain further biological information on these intriguing structures formed on the cell surface, the same sample was transferred to a synchrotron facility for the elemental mapping using SR µXRF (Figure 3E), supported by scanning transmission X-ray microscopy (STXM) (Figure 3F) for simultaneous mapping of atomic number by absorption contrast sensing.^68^ The spectral maps of primarily light elements have been generated from the spectral data and their fits, where we show an example from the site marked with an asterisk. The color-coded theoretical spectra for each of the measured elements fit perfectly to the raw data (in black) with a high signal to noise ratio (S/N) due to the high detection sensitivity of the technique. The results confirm not only a high TiO_2_ content within highly spherical composites, as evidenced by locally elevated concentrations of O (K-shell emission spectra) and Ti (much weaker L-shell emission spectra), but also a higher content of C, N, Mg and, to some extent, Na (all K-shell emission spectra) at the same locations. This again confirms the extensive physical affinity of biomolecules, such as lipids or cholesterol, shown previously^39^ and positive ions to the surface of the TiO_2_ NTs as elaborated in the first results section. It further validates and signifies our recent findings performed on the same nano-bio system using a combination of omics, *in vitro* measurements and full-atom *in silico* simulations, which reported strong interaction of amine and phosphate groups of the disordered lipids with the TiO_2_ NTs surface.^39^ Extensive biomolecule binding, forming the so-called lipoprotein corona as clearly demonstrated by the ultra-high resolution imaging (Figure S4), can significantly disrupt cellular homeostasis by altering protein function, affecting cell signaling and trafficking machinery,^73^ all of which can potentially lead to immune activation and long-term cellular stress. The local increase in Na^+^ adsorption may be due to the high surface/volume characteristic of anatase TiO_2_ NTs, allowing Na^+^ to intercalate into the surface and subsurface layer.^74^ The most interesting observation is the deviation of the distribution of plausibly extracellular Na^+^ from the rest of the elemental maps, indicating at least partially a different binding mechanism. Using correlated µXRF and SEM, we observe complete co-localization of the highly elevated Na^+^ sites with the distinct filamentous structures formed by cells bridging the TiO_2_-bio composites (Figure 3E, bottom image, purple arrows). Good correlation of chemical maps with distinct physical properties not only highlights a fundamental link between composition and structure, but also suggests a potential functional relationship. The preferential binding is likely due to the high concentration of negatively charged biomolecules within the filamentous structure. Among these, heparan sulfate (HS) chains stand out, as they possess the highest negative charge density of any known biological macromolecule and are commonly found on cell surfaces.^75^ These molecules also play a central role as mediators in inflammatory processes,^76^ suggesting local inflammatory activity on TiO_2_-bio composites bridged with these filamentous structures. In addition, the sequestration of both intrinsic and/or extracellular Na^+^ and Mg^2+^ may result in alteration of ion homeostasis, disruption of normal associated cellular functions and induction of oxidative stress.

To further elucidate the intricate morpho-functional properties of the studied nano-bio interface at the nanoscale, we correlated multimodal CLSM with SR µXRF and ultra-high resolution helium ion microscopy (HIM), performed for the first time (Figure 4).

**Figure 4.**
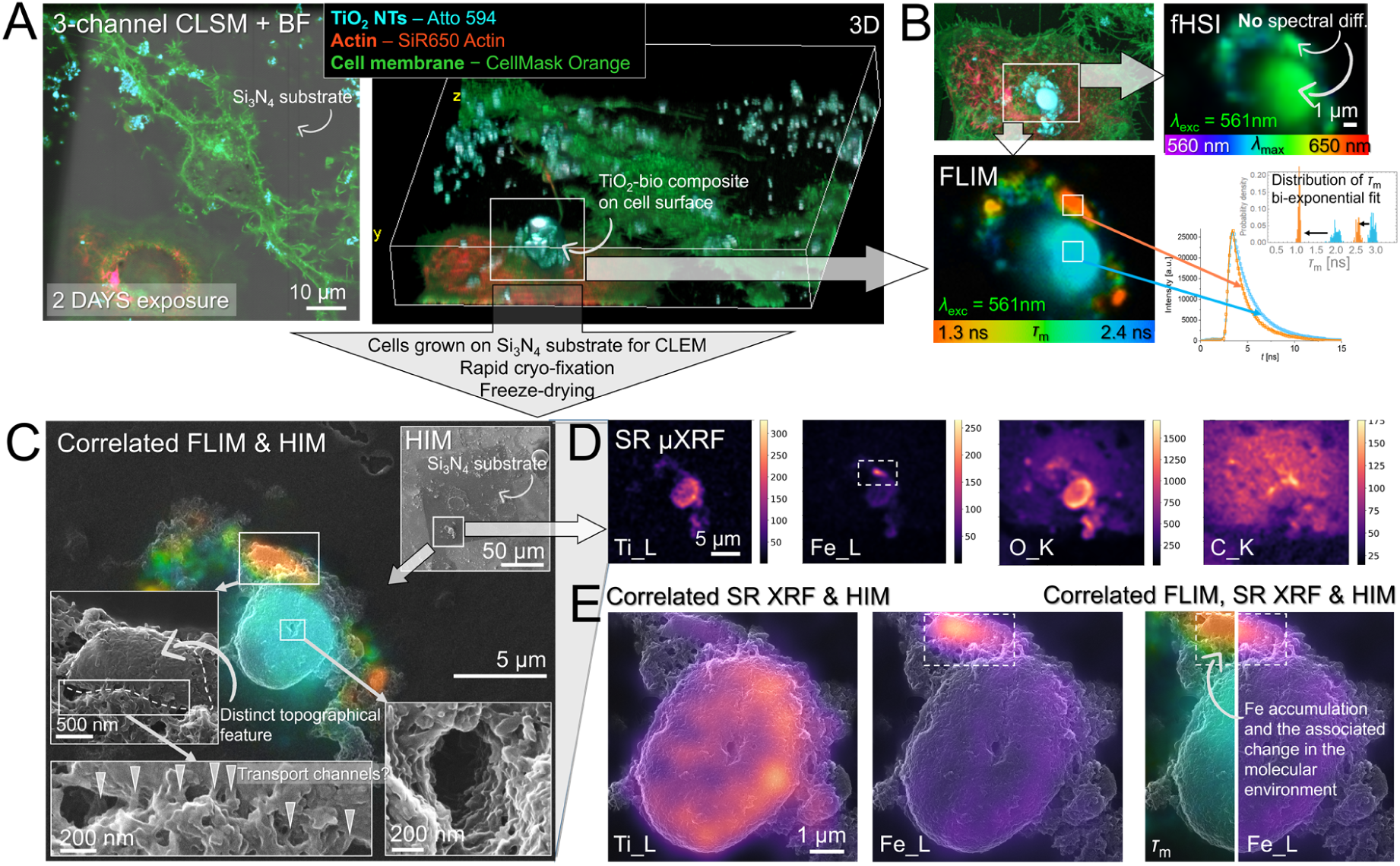
Novel correlated high-resolution FLIM, SR µXRF and HIM providing in-depth morpho-functional assessment of inflammatory TiO_2_-bio cell-excreted composites immobilized on the surface of lung epithelial LA4 cells. A) Cells grown on thin Si_3_N_4_ transparent substrate (in gray, BF image) with labeled cell membranes (in green), actin (in red) and nanoparticles (in light blue) acquired with 3-channel CLSM using 60X water immersion objective. 3D CLSM revealing size, shape and topography of a large, few micron-sized TiO_2_-bio composite formed on the cell surface (marked with an arrow). B) fHSI and FLIM imaging and analysis of the investigated structure uncovering no particular spectral, but significant fluorescence lifetime contrasts, respectively (see the charts on the right). C) Correlated FLIM and HIM exhibiting co-localization of distinct fluorescence lifetime map (in orange) and nanoscale topographical features marked in the left inset image. Further ultra-high magnification inset images on the bottom uncover intriguing morpho-functional features, from the apparent multiple transport channels into the local site with an accumulated Fe (marked with arrows) to the void or tunnel formed in the center of the TiO_2_-bio composite extending towards the plasma membrane. D) SR µXRF elemental maps of metals and biomolecules distributed locally and more evenly throughout the investigated cellular region. E) Changes in physical properties of the molecular environment on the nano-bio interface plausibly caused by high Fe accumulation shown by correlated FLIM, SR µXRF and HIM, the correlative microscopy approach applied for the first time.

By targeted fluorescence labelling prior to live cell imaging of lung epithelial LA4 cells, that had been exposed to TiO_2_ NTs for a period of 2 days, we were able to perform a detailed characterization of highly specific nano-bio structures formed on the cell surface (Figure 4A-B). Again, a large, condensed TiO₂ NTs-biostructure, up to a few microns in size, with an approximately spherical shape, was observed and imaged in detail in the multichannel 2D and 3D (Figure 4A). Moreover, we performed additional FLIM and fHSI imaging (Figure 4B), which allowed us to investigate the potential heterogeneity of the molecular environment in the immediate vicinity of the reporting fluorophores attached to the surface of TiO_2_ NTs. While fHSI showed no significant spectral variation across the composites, indicating the absence of any particular local molecular and physical changes in the immediate nm vicinity, FLIM, revealed significant deviations in the fluorescence lifetimes (*τ*_m_) within the same locations, as indicated by the orange and blue rectangles. This may be indicative of different degrees of dye aggregation/self-quenching^77^ due to the different surrounding molecular environment, but a more in-depth investigation using an expanded set of complementary techniques, is required to better understand the cause.

To elucidate the underlying cause of the observations, the sample, grown on Si_3_N_4_ substrate was cryo-fixated immediately after fHSI/FLIM imaging and transferred to successive correlated high-vacuum SR µXRF and ultra-high resolution HIM for the most accurate chemical and morphological analysis, respectively. Chemical speciation and detailed surface examination provided new insights that now better explain the differences in the molecular environment observed in living cells (Figure 4C-E). Using extremely sensitive, sub-micron resolution SR µXRF, we revealed the presence of localized, highly concentrated iron (Fe) (indicated by the dashed rectangle), as opposed to more widely distributed titanium (Ti), and in particular intrinsic carbon (C) and oxygen (O). By superimposing the data from all three complementary techniques (Figure 4E, on the right), we observe a remarkable co-localization of the concentrated Fe (in pink), reduced fluorescence lifetime (in orange), supported by the distinct morphological and topographical feature within the same region provided by ultra-high resolution HIM (Figure 4C, left inset). With our novel correlative microscopy approach, we not only identified a close relationship between FLIM footprint, Fe content and distinct surface morphology, but also gained a better morpho-functional assessment of the observed nano-bio structure to better elucidate possible causes and consequent biological effects, as discusses here.

The site of highly localized Fe, accompanied by the numerous distinct morphological features at the outer edge, suggesting possible biological transport channels in the form of tunneling nanotubes (TNTs)^78^ with an average diameter of 50 nm (Figure 4C, marked with arrows), implies the biological cause of its accumulation through the active cell secretion. This also implies the nanoparticle-induced alteration of iron homeostasis and the subsequent cell-protective mechanism, through the initiated cellular iron efflux, plausibly via the major iron transporter ferroportin (Fpn).^79^ This mechanism opts to mitigate iron-induced reactive oxygen species (ROS) and pro-inflammatory cytokine production and further iron-mediated inflammation that can damage DNA, proteins and lipids.^80^ The remaining question is, why the accumulation in the TiO_2_- bio composite? These large TiO_2_-rich extracellular vesicles may serve as extracellular containers for the other waste molecules, where their high absorptivity at the metal oxide surface, such as ferrous iron,^81^ may also play an important role.

To better understand the physical aspects of the interaction, we need to understand the corresponding reduced fluorescence lifetime of the biological molecules present at the nano-bio interface. The proximity and/or adhesion of Fe particles to the TiO_2_ surface could potentially induce a phenomenon of localized surface plasmon resonance (LSPR)^82^ resulting from the laser-induced collective oscillations of surface electrons in a conduction band,^83^ leading to a shorter fluorescence lifetime.^84^ Conversely, Fe could interact with nearby lipids and proteins enveloping TiO_2_, catalyzing lipid peroxidation,^85^ inducing lipid bilayer disruption through an increase in water permeability and thus an increase in membrane polarity,^86^ resulting in a shortening of the dye lifetime.

The implementation of correlative ultra-high resolution and large depth-of-field HIM revealed another intriguing feature at the center of the object, resembling a ‘wormhole’ (inset on the right). The more than 1 µm deep void, or tunnel, was formed during the assembly of a TiO_2_-bio composite and could potentially be used for the diffusion and selective matter exchange between the inside and outside of the cell. This type of nano-bio interface assembly with the revealed nm features has never been demonstrated before and could be an interesting subject for further studies.

Finally, we uncover one of the potentially crucial initiating modes of action in the inflammatory cell response to TiO_2_ NTs. Using high magnification SEM, we have observed and characterized a thin fibrous network of biological material that has developed locally over a large TiO_2_-bio composite immobilized on the cell surface (Figure 5D). The organization and the thickness of the fibers of a few tens of nanometers are reminiscent of fibrin.^87,88^ Considering that fibrin has been found to be excessively deposited in the early stages of acute intra-alveolar lung injury,^89,90^ plausibly accompanied by the secretion of fibrinogen by lung epithelial cells,^91^ the possible presence of fibrin on our studied nano-bio interface could have significant biological implications for the local pro-inflammatory stimuli accompanied by the secretion of cytokines, such as tumor necrosis factor (TNFα).^92^ To better elucidate the possible TNFα secretion-induced fibrin fiber formation, we performed an additional TiO_2_ NTs exposure experiment on the co-culture of lung epithelial LA4 and murine alveolar macrophages MH-S, which are known to release TNFα after exposure to metal oxide nanoparticles^93^ or high aspect ratio nanomaterials^94^ such as ours. Our aim was to determine whether the presence of MH-S increased the formation of fibrous structures locally on the TiO_2_ NTs. The results reveal a significantly higher presence of fibrous structures on nearly all TiO₂ NTs immobilized on the lung epithelial cell surface in the co-culture, compared to the partial—yet still substantial—formation observed in the monoculture, primarily over the larger composites (Figure S5). We also observe an extensive apical MH-S scavenging and uptake of TiO2 NTs (Figure S6), where rounded macrophages with phagocytic protrusions capture and trap the nanoparticles, identified by the white image contrast.

**Figure 5.**
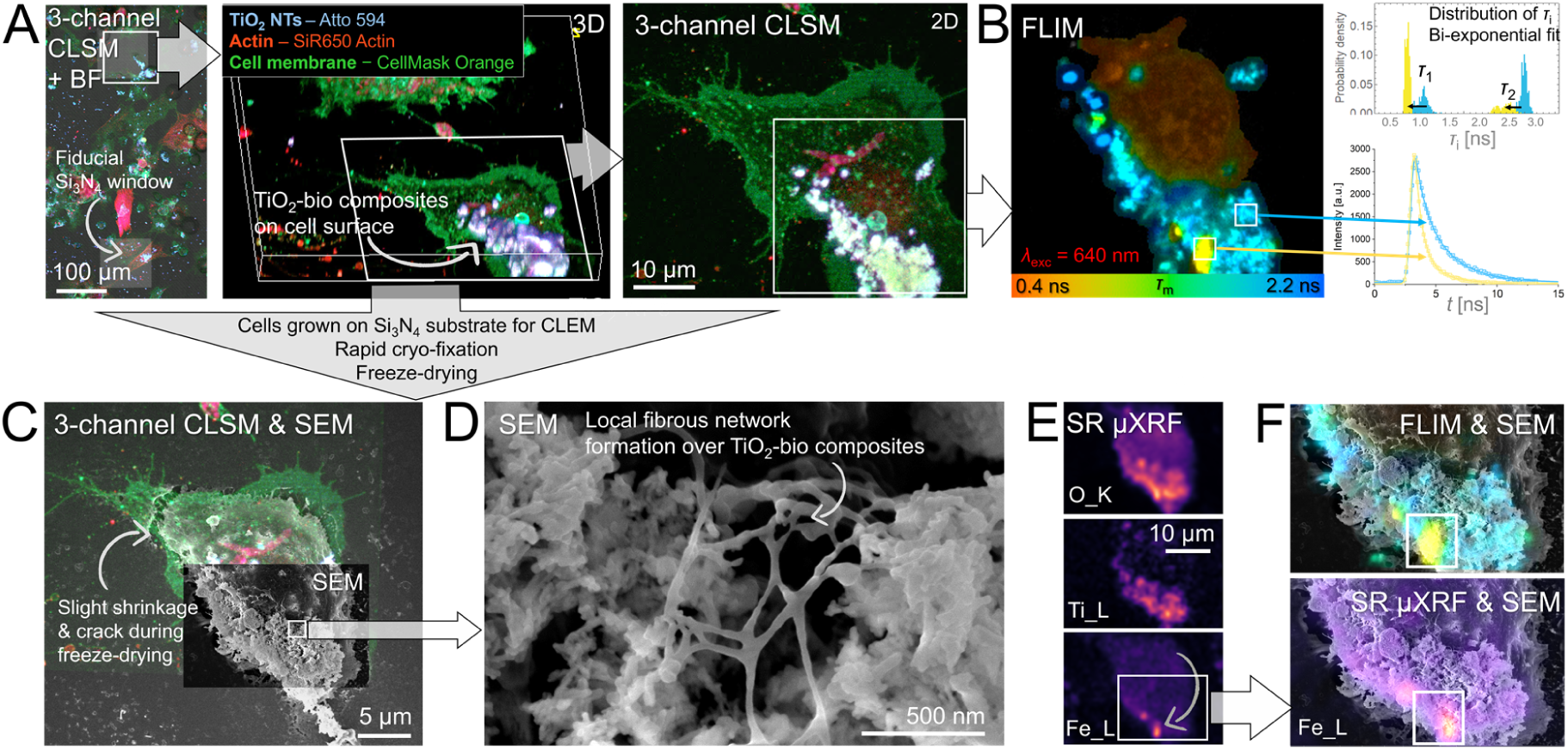
Novel high-resolution CLEM-µXRF confirming Fe-induced changes in the physical properties of the surrounding biological environment and uncovering the plausible inflammatory local formation of fibrous network resembling fibrin matrix formed over TiO_2_-bio composites immobilized on the plasma membrane of lung epithelial LA-4 cells. A) Cells grown on thin Si_3_N_4_ substrates (in gray) with labeled cell membranes (in green), actin (in red) and nanoparticles (in blue) acquired with 3-channel CLSM using 10X and 60X objective magnification. High-magnification 2D and 3D CLSM reveal size, shape and topography of a large, more than ten micron-sized TiO_2_-bio composite arrested on the surface (marked with an arrow). B) FLIM imaging (on the left) and analysis (on the right) of the investigated composite revealing significant contrast in fluorescence lifetime between local regions (from yellow to blue). C) Correlated CLSM and SEM showing slight shrinkage and local cracks near the cell boundary after rapid freeze-drying but, most importantly, preserving the investigated structures on cellular surface. D) High-magnification SEM uncovering local formation and the morphology of fibrous network formed over the TiO_2_-bio composites on the cellular surface, with its inflammatory implications further examined in the supporting Figures S4-5. E) SR µXRF elemental maps of metals and oxygen distributed throughout the investigated cell. F) Confirmation of Fe-induced local physical changes on the nano-bio interface (inset regions) measured by correlated FLIM and SR µXRF represented on the top of SEM data.

To confirm that the fibers consist of inflammation-mediating fibrin molecules, we performed another correlative experiment by introducing labeled fibrinogen to the living cells exposed to TiO_2_ NTs, allowing us to assess potential co-localization with the fibrous structures using the following high-resolution HIM (Figure S7). Despite the much lower resolution of the confocal microscopy images and the slight sample displacement during sample preparation, preventing the precise registration and overlap with HIM, the results still show a good co-localization of the fibrous network with the labelled fibrinogen, further supporting our hypothesis. To further explore the potential of long-term inflammation and fibrogenesis in the development of such structures over time, further studies are needed.

In addition to the fibrous structures observed by high magnification SEM and HIM, we provide a correlative characterization of the same TiO_2_-bio composites using multimodal live cell 3- channel CLSM and FLIM (Figure 5A-B) and SR µXRF (Figure 5E-F). FLIM revealed a high degree of heterogeneity in the fluorescence lifetimes of Atto 594 dye attached to TiO_2_ NTs. The average lifetime *τ*_m_, calculated from a bi-exponential fit, was approximately half shorter in localized regions (in yellow) than in the rest parts of the sample (in green to blue). Correlated SR µXRF again showed a co-localization between the short fluorescence lifetime and the site with high Fe accumulation (Figure 5E-F) as reported in Figure 4, suggesting a similar underlying mechanism.

## Conclusion

In this study, we highlight the importance of implementing novel multimodal and multiscale extended CLEM, to elucidate the intricate interactions between biological matter and nanoparticles. In particular, we present different response mechanisms of lung epithelial cells, a subject of persistent exposure to particulate matter, to exemplary high aspect ratio TiO_2_ nanotubes with known subacute and even chronic inflammatory inducing effects. The study provides a comprehensive morpho-functional assessment of the nanoparticle-cell interface from the micron down to the nanoscale with the implications for lung epithelial toxicity through the uncovered possible initiating modes of action as schematically summarized in Figure 6. With the novel correlative microscopy approach and in-depth analysis, we were able to advance the understanding of the recently reported TiO_2_-induced mechanism of chronic inflammation^39^ and genotoxicity^41^ by better elucidating several possible cell response mechanisms to the disruptive nature of this particular nanoparticle with different inflammatory and anti-inflammatory outcomes:

`o` Nearly spherical TiO_2_ NTs-bio composites with extensive DNA binding immobilized on the cell surface, similar to immobile apoptotic bodies, may result in defects in apoptotic cell clearance. This may lead to secondary necrosis, loss of membrane integrity and release of immunostimulatory contents through the observed plasma membrane lesions.
`o` The formation of distinct filamentous structures bridging TiO_2_-bio composites, perfectly co-localized with high Na^+^ accumulation, indicating a localized, highly negative charged surface of plausible heparan sulfate, a key mediator in cell inflammation.
`o` Nanoparticle-induced alteration of iron homeostasis and balance of other TiO_2_-adherent cellular compounds/biomolecules, followed by the controlled release of an excess iron or other exchange compounds through transport channels, suggesting a cell-protective mechanism.
`o` The localized formation of a fibrin-like fibrous network over TiO_2_-bio composites indicating on the local pro-inflammatory stimuli accompanied by plausible secretion of fibrinogen and TNFα cytokines.

**Figure 6.**
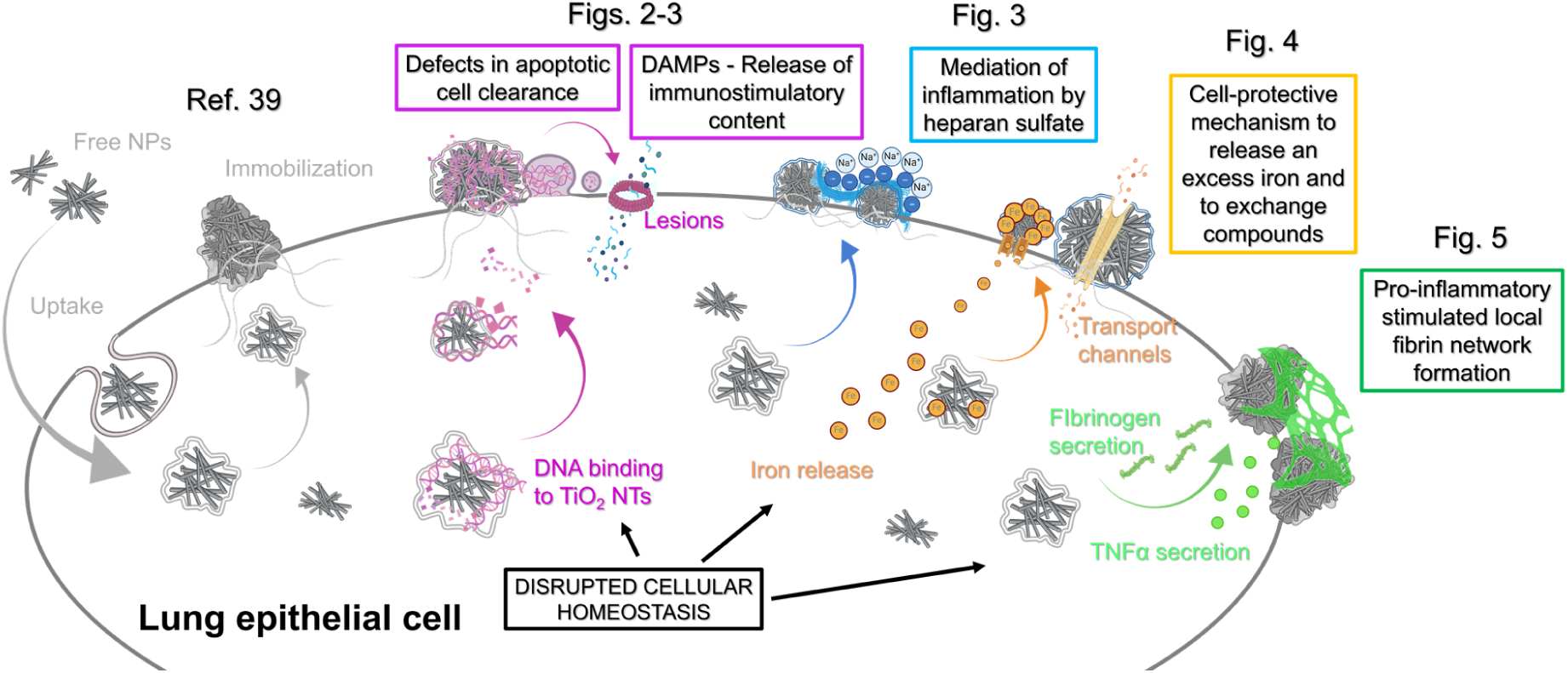
The schematic overview of the possible modes of action of the cell response to the disruptive nature of TiO_2_ NTs, toxic to lung epithelial cells, with different inflammatory and anti-inflammatory outcomes (colored from pink to green), observed at the nano-bio interface.

By implementing such a multimodal and multiscale correlative microscopy approach for the first time, we were able to study and characterize the initial cellular responses to exposed nanoparticles at such a small scale. These responses may play a critical role in the body’s initial immune response and the development of inflammation.

This methodology provides a strong basis for advancing research into the toxicity of nanoparticles and their potential impact on human health. By providing a robust framework, it enables further exploration and refinement of approaches to toxicity assessment. However, to fully realize its potential and guide future research, it is essential to address certain limitations. Key challenges include the limited high-throughput imaging required for improved statistics, together with faster acquisition to minimize sample displacement, and the need for optimization and better control of all physical variables throughout sample preparation to preserve nanoscale features throughout the sample. Both of these challenges point to better automation, which is discussed in Supplementary Comment #2 in the Supporting Information Appendix. This will lead to critical improvements in the future for more precise 2D/3D visualization, evaluation and deeper insights into nano-bio interactions and the mechanisms behind them, paving the way for more accurate and reliable interpretations of how different nanoparticles and their properties affect relevant biological systems, important for improved safety assessment.

## Experimental section

### Material

The murine epithelial lung tissue cell line (LA-4, ATCC CCL-196), murine alveolar lung macrophage cell line (MH-S; ATCC, CRL-2019), F-12K medium (Gibco), Fetal bovine serum (ATCC), 1% Penicillin-Streptomycin (Sigma), 1% non-essential amino acids, L-glutamine, beta-mercaptoethanol (Gibco), Phosphate buffer saline (PBS), Live Cell Imaging Solution (LCIS, Invitrogen); Titanium dioxide nanotubes (TiO_2_ NTs) in anatase form synthesized in-house^95^; μ- Slide 8-well (Ibidi), Silicon Nitride Si_3_N_4_ Support Film (PELCO, 21509CL, Ted Pella), TEM formvar/carbon film on Au Gilder 200 F1 finder grids (FCF200F1-AU-50, EMS), Ammonium acetate (Sigma-Aldrich), PTFE coated high precision and ultrafine tweezers (72919-3SATe, EMS), propane transfer system for plunge freezer (37015, Electron Microscopy Sciences); AlexaFluor 647 (Thermo Fischer Scientific), Atto 594 (ATTO-TEC), CellMask Orange (Invitrogen), SiR-actin (Spirochrome), Draq5 (Invitrogen), MitoTracker™ Orange CMTMRos (Invitrogen), PSM-39 (in-house).

### Cell culture

The LA-4 murine epithelial lung tissue cells were cultured in 75 cm² TPP cell culture flasks. They were maintained in a controlled environment at 37°C with 5% CO₂ and humidity using F-12K medium supplemented with 15% fetal bovine serum (FBS), 1% penicillin-streptomycin, 1% non-essential amino acids and 2 mM L-glutamine The MH-S alveolar macrophage cells were cultured in RPMI 1640 medium (Gibco), supplemented with 10 % FBS, 1% penicillin-streptomycin, 1% non-essential amino acids, 2 mM L-glutamine and 0.05 mM beta-mercaptoethanol (Gibco). Upon reaching 70-90% confluency at the appropriate passage, the LA-4 cells were seeded onto specially designed imaging holders customized for correlative microscopy. To enable both, reflection and transmission imaging modes, thin Silicon Nitride (Si_3_N_4_) and formvar/carbon on Au Gilder 200 F1 finder grid support films were used. For co-culture experiments, LA-4 and MH-S were grown in separate dishes up to 70-90% confluency and mixed together by the ratio LA- 4:MH-S, 40:1. After 24h incubation, the co-cultures were exposed to nanoparticles for the desired amount of time, prior microscopy. Growth medium in the co-culture was an equal mixture of F- 12K and RPMI 1640.

### Nanoparticles

TiO₂ nanoparticles have been selected for investigation in this study due to the potential risks to human health associated with inhalation or oral exposure.^96,97^ In case of their high specific surface area, recent studies have demonstrated the potential for these nanoparticles to elicit an inflammatory response ^39,40^ and to possess carcinogenic properties.^41,98^ TiO₂ nanotubes (TiO₂ NTs) in anatase form synthesized in-house with the protocol described in ^95^ exhibited a diameter range of 6 to 11 nm, a mean length of 100-500 nm, and a BET surface area of 150 m^2^ g^-1^.^40^ In order to facilitate enhanced visualization and monitoring of nanomaterial interactions and their impact on model lung epithelium at the sub-micron scale, the TiO₂ NTs were functionalized with super-resolution compatible fluorescent probes, AlexaFluor 647 and Atto 594, according to the protocol avoiding the introduction of artefact as described in ^48^. Functionalized and filtered nanoparticles suspension was stored in a 100-fold diluted bicarbonate buffer (5 mOsm L^-1^) prior administration to cells.

### Sample preparation for live-cell confocal fluorescence microscopy

After reaching 70-90% confluency in culture flasks, LA-4 cells were reseeded directly onto UV-sterilized Si_3_N_4_ or formvar/carbon Au finder grid substrates/holders placed within the chamber of a μ-Slide 8-well. The volume of the added cell suspension prepared at a concentration of a few 10⁵ cells/ml was 20 μl and resulted in approximately 5,000 cells per holder with a diameter of 3 mm. Following a four-hour period during which sufficient cell attachment to the substrate had occurred, cell growth media was added to each chamber in order to cover the surface and wet the entire holder, thus preventing any evaporation during the course of the measurements. Once the desired confluency of approximately 50% had been reached (typically within two days), gently sonicated (Bransonic ultrasonic cleaner, Branson 2510 EMT) nanoparticle suspension and carefully rinsed onto the cell monolayer at an average calculated surface dose of 5:1 (*S*_nanomaterial_:*S*_cells_) and total 3 μg/cm². It presents a meaningful concentration according to the lifetime occupational exposure and the safety recommendations as further discussed in Supplementary Comment #1. The surface dose was achieved by pouring *V* = 5 μl of a suspension of nanoparticles (*c* = 0.5 mg/ml) into the chambers of 1 cm² size filled with the cell growth media (*V* = 300 μl). To put into perspective, the surface dose, which is the most important determinant in the nanomaterial toxicity in lungs,^99^ was used to be equivalent to exposure over a period of 45 working days, based on the 8-hour time-weighted average occupational exposure limit for TiO_2_ (6.0 mg/m^3^ TiO_2_) established by Danish Regulations.^100^ The cells were exposed to nanoparticles for the desired period of time, which was one to three days, prior to live-cell imaging and following cryopreservation for high-vacuum imaging.

Before live-cell imaging, cells were stained with different highly-specific fluorescent dyes to visualize different cell compartments potentially interacting or responding to exposed nanoparticles. Staining was performed according to the manufacturer’s recommendations to achieve sufficient fluorescent signal while preserving cellular functions. SiR Actin dye in concentration *c* = 400 nM was used for 4 h to stain F-actin. Draq 5 dye in concentration *c* = 5 µM was used for 15 min to stain double-stranded DNA. CellMask Orange in 1000X diluted stock concentration was used for 10 min to stain cell plasma membrane. Mitotracker Orange in concentration 500 nM was used for 15 min to stain mitochondria. All staining solutions were removed and sample was rinsed a few times with LCIS prior measurements performed on the microscope inside the stage top incubator with a temperature, gas and humidity control (H301- MINI, Okolab). Before measurements, the implemented holders with the adhered cells were flipped to enable imaging with small working distance (high numerical aperture) objective. Sterilized polystyrene microspheres of 20 µm in size were employed as volume spacers in order to prevent any compression of the sample.

### Sample preparation for high-vacuum electron and ion microscopy

Immediately after live-cell imaging, sample was carefully transferred from the measuring μ-Slide 8-well into the 150 mM ammonium acetate washing solution to eliminate buffer salt crystals, which otherwise interfere with both structural and chemical analysis performed with high-vacuum techniques.^71^ After few seconds of immersion in the washing solution for three times, an excess of water was removed by careful blotting the sides of the sample holder with a lint-free paper. Immediately after blotting, samples were rapidly cryofixated without chemical fixation, to preserve the integrity and morphology of cellular structures and interacting nano-bio interface on the nanoscale, using plunge freezing in a liquid propane to prevent the formation of crystalline ice^101^ and stored in a cryo bank until the final drying procedure in a freeze dryer (Coolsafe 100-9 Pro). Propane was prepared in a liquid nitrogen-cooled chamber with the plunge freezer transfer system. To carefully handle with the fragile specimens, high precision cryogenic and non-sticking PTFE tweezers were used. In case of chemical fixation, ammonium acetate washing solution was changed with 4% paraformaldehyde (PFA) and 2% glutaraldehyde (GA) solution. Samples were fixated for 15 minutes at room temperature before blotting and rapid cryofixation.

### Confocal laser scanning microscopy (CLSM)

The experiments were conducted on a custom-built inverted microscope (Olympus IX83, Abberior Instruments). Imaging was conducted with Olympus 10x (*NA* = 0.3) and 60x (*NA* = 1.2) water immersion objectives, utilizing two pulsed ps lasers in conjunction with a fast-gating system, which was controlled by an FPGA unit. In order to facilitate the simultaneous detection of the fluorescence emitted by different fluorophores, two pulsed diode lasers were employed, with wavelengths of 561 nm and 640 nm, respectively, and a pulse length of 120 ps and repetition rate of 80 MHz. Fluorescence was detected in a descanned mode using an additional pinhole to reduce out-of-focus light and avalanche photodiodes (APD, SPCM-AQRH, Excelitas) with a photon detection efficiency (PDE) exceeding 50% across the entire visible spectrum. Fluorescence was collected from each scanning voxel using a dwell time of 10 µs and detected within the spectral windows λ = 580 - 625 nm and λ = 650 - 720 nm, utilizing dichroics and band-pass filters (both Semrock). For more comprehensive illustration of the optical setup, please refer to the schematics in Figure S8. Supporting high-resolution measurements on live cells were conducted with the integrated customized super-resolution microscopy (STED), using depletion laser wavelength 775 nm, pulse length of 1.2 ns and the power of 200 mW in the sample plane.

### Fluorescence lifetime imaging microscopy (FLIM) and fluorescence hyperspectral imaging (fHSI)

The microscope system was upgraded with the 16-channel Multi-Wavelength Photo-Multiplier Detectors (PML-16 GaAsP, Hamamatsu), a multi-dimensional Time Correlated Single Photon Counting (TCSPC) detection system and a DCC-100 detector controller card (all manufactured by Becker&Hickl), all communicated through fast ps FPGA electronics, enabling the simultaneous detection of signals across up to 16 spectral channels and detection of their corresponding fluorescence lifetime decays.^102^ For fluorescence lifetime imaging, the signals collected on the 16 GaAsP detectors that arrived in the same time interval were summed and the FLIM decay curves obtained in each pixel of the image analyzed using the SPCImage5.0 software (Becker&Hickl) as done before.^103,104^ The analysis of FLIM decay curves was conducted using a two-component, double exponential fitting method convoluted with the instrument response function (IRF), which constrains the achievable resolution to its 200 ps FWHM. The pixel exposure time (100 µs) and image binning (3×3) were set to obtain a sufficient peak value of the decay curves of at least 1000 to eligibly resolve fitting with a double-exponential model:

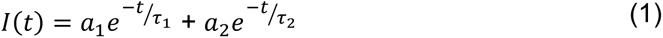

with the lifetime values of the fast and slow decay components, *τ*_1_ and *τ*_2_, and the corresponding intensity coefficients, *a*_1_ and *a*_2_. The fitted FLIM decay curves from each pixel were color-coded according to the average fluorescence lifetime *τ*_m_ = *a*_1_*τ*_1_ + *a*_2_*τ*_2_, with the fastest decays shown in red and longer decays in blue. Furthermore, fHSI was conducted utilizing 16 spectral channels within the wide spectral window spanning from 560 nm to 760 nm, as determined by the diffraction grating positioning within the spectrograph (PML-Spec, Becker&Hickl). The high quantum efficiency of approximately 50% across the whole visual spectrum was achieved by GaAsP PMT detectors. The total number of photon counts collected on TCSPC for each pixel was calculated for each spectral channel and represented in the spectral curve. To avoid partial blocking of photons on the notch filters in our de-scanned detection setup which distorts the acquired fluorescence spectra, an optical bypass for collected photons was developed using a combination of acousto-optic tunable filters (AOTF, Opto-electronic) (Figure S1). The hyperspectral data were fitted by a suitable spectral model, namely the most convenient empirical log-normal function due to its numerical stability during optimization^69^:

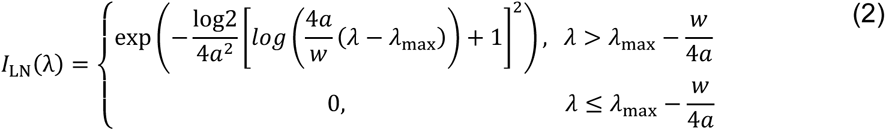

with peak position (*λ*_max_), approximate full width at half-maximum (FWHM, *w*) and asymmetry (*a*). The developed model was demonstrated to be capable of resolving changes in the spectra as small as 1 nm.^69^ Pixel binning of 3×3 was employed prior to spectral fitting, increasing sensitivity at the cost of image resolution.

### Scanning electron microscopy with energy-dispersive X-ray spectroscopy (SEM-EDS)

High-resolution imaging and chemical analysis were conducted in the FEI Helios Nanolab 650 scanning electron microscope (SEM) utilizing energy-dispersive X-ray spectroscopy (EDS). To prevent the charging effects of a highly insulating biological specimen during electron irradiation, all samples were coated with an approximately 10 nm carbon layer. High-resolution and high surface sensitivity imaging was performed with low electron acceleration voltage (2 keV), low electron current (100 pA) and under high chamber vacuum (10^-6^ hPa). The EDS spectra were collected with higher electron acceleration voltage (15 kV) and electron current (200 pA). Under these conditions, an energy resolution of 145 eV at the Mn Kα line and an elemental sensitivity of approximately 0.01 wt% (X-Max SDD, Oxford Instruments) were achieved. The weight percentage (wt%) of each individual element was calculated and subsequently employed for further analysis, which entailed the measurement of the differences in the wt% ratios of the studied elements between individual regions.

### Helium Ion Microsopy (HIM)

High-resolution images at the nanoscale were obtained using a helium ion microscope (Orion NanoFab, Zeiss).^105^ For imaging purposes, He atoms are ionized in the GFIS source. The source is characterized by a small virtual source size (0.25mm), a high reduced brightness (1-4 10^9^ Am^- 2^sr^-1^V^-1^) and a very small energy spread of less than 1 eV. Subsequently the beam is focused, shaped and scanned over the sample surface using electrostatic lenses, quadrupoles, and octapoles. The resulting focused ion beam^106^ has a diameter of 0.5 nm and enables the recording of images with a large depth of focus.^107^ Prior to imaging, the freeze-dried samples grown on Si_3_N_4_ substrate were affixed with carbon tape to the standard pins compatible with a multi-pin specimen mount. Insulating samples were spared of conventional thin layer conductive coating due to efficient charge compensation within HIM instrument using the flood gun^108^, thus preventing any surface modification at the nanoscale studied. Imaging was conducted through the detection of secondary electrons (SE1) emitted from the few nm surface layer of the sample. The following experimental parameters were employed: helium ion energy (30 keV), ion current (0.5 pA) and chamber vacuum (2×10^-7^ hPa). The field of view (FoV) of the acquired images ranged from less-magnified, 175 µm by 175 µm required for registration, to highly-magnified, 0.7 µm by 0.7 µm, with a minimal pixel step size of 0.7 nm and typical dwell time 20 µs at each pixel.

### Synchrotron Micro-X-ray Fluorescence (SR µ-XRF)

The X-ray fluorescence (XRF) data were collected at the TwinMic beamline of Elettra Sincrotrone Trieste, located in Trieste, Italy.^109^ The TwinMic microscope was operated in scanning transmission mode, whereby the sample was raster scanned across an incoming perpendicular X-ray beam, delivered by a zone plate diffractive optics. During the scanning process, a rapid readout charge-coupled device (CCD) camera (DV 860, Andor Technology) collected the transmitted X-ray photons, resulting in the generation of absorption and differential phase contrast images.^110^ Simultaneously, the emitted X-ray fluorescence (XRF) photons were collected by eight silicon drift detectors (SDD) situated in front of the sample at an angle of 20 degrees with respect to the sample plane.^111^ This results in the simultaneous acquisition of both, the morphology of the sample and the distribution of the elements.

In order to achieve optimal excitation of Mg, Na, O, Ti and C, while also obtaining sub-micron spatial resolution, a monochromatic energy of 1.5 keV was selected for the experiment. The specimens were scanned at a step size of 500 nm with a probe size of 600 nm in diameter, delivered by an Au zone plate of 600 µm diameter and 50 nm outermost zone width. An acquisition time of 50 ms pixel⁻¹ and 4 s pixel⁻¹ were employed for the CCD and SDD detector systems, respectively. The acquired XRF spectra were processed using the PyMCA software.^112^

## Supporting Information

Additional high resolution and correlative microscopy results of the nano-bio interface supporting the investigated possible modes of action leading to nanoparticle induced lung epithelial toxicity.

## Acknowledgements

This work was supported by Helmholtz European Partnering project CROSSING (Grant No: PIE- 0007), EU H2020 project no. 824096 “RADIATE”, the Slovenian research and innovation agency (ARIS) programs P1-0060 (Experimental biophysics of complex systems), P1-0112, I0-0005 and project N1-0090. The authors thank Prof. Dr. Katarina Vogel Mikuš of the Biotechnical Faculty (University of Ljubljana) for her invaluable assistance with the freeze-drying of samples, Dr. Valentina Bonanni of the Elettra Sincrotrone (Trieste, Italy) for help with the SR µXRF measurements, Prof. Dr. Janez Štrancar of the Jožef Stefan Institute (Ljubljana, Slovenia) for initiating the scientific field in the Laboratory of Biophysics, Dr. Tilen Koklič for fruitful discussions, Dr. Iztok Urbančič for the hyperspectral analysis software support, DR. Stane Pajk for the synthesis of lipid droplet dye, Dr. Polona Umek for synthesis of TiO_2_ nanoparticles and Dr. Hana Kokot for fluorescence labeling of TiO_2_ nanoparticles (all of the Jožef Stefan Institute).

## Author contributions

R.P. conceptualized and designed the analysis; R.P., A.K., L.P. and A.G. performed the experiments; R.P., L.H. and A.G. collected the data; R.P. performed the analysis and data visualization; R.P., G.H. and A.G. contributed analysis tools; R.P. wrote the paper; R.P., L.P., G.H., A.G. and P.P. reviewed and edited the paper; P.P. acquired funding.

## Notes

### Competing Interest Statement

The authors have declared no competing interest.

### Summary of Updates

We provide additional high resolution and correlative microscopy results of the nano-bio interface (new Supporting Figures S1-S7) accompanied by the more detailed biological and physical interpretation of the investigated possible modes of action leading to lung epithelial toxicity, as shown in the new schematic figure (Figure 7).

## References

(1) Correlative Light and Electron Microscopy IV, Volume 162 - 1st Edition, In Methods in Cell Biology, Reichert, T. M.; Verkade, P., Eds.; Academic Press, 2021; pp 2–430.

(2) Ando, T.; Bhamidimarri, S. P.; Brending, N.; Colin-York, H.; Collinson, L.; Jonge, N. D.; Pablo, P. J. de; Debroye, E.; Eggeling, C.; Franck, C.; Fritzsche, M.; Gerritsen, H.; Giepmans, B. N. G.; Grunewald, K.; Hofkens, J.; Hoogenboom, J. P.; Janssen, K. P. F.; Kaufmann, R.; Klumperman, J.; Kurniawan, N., et al. The 2018 Correlative Microscopy Techniques Roadmap. J. Phys. D: Appl. Phys. 2018, 51 (44), 443001.

(3) Datta, R.; Heaster, T. M.; Sharick, J. T.; Gillette, A. A.; Skala, M. C. Fluorescence Lifetime Imaging Microscopy: Fundamentals and Advances in Instrumentation, Analysis, and Applications. J Biomed Opt 2020, 25 (7), 1–43.

(4) Sekar, R. B.; Periasamy, A. Fluorescence Resonance Energy Transfer (FRET) Microscopy Imaging of Live Cell Protein Localizations. J Cell Biol 2003, 160 (5), 629–633.

(5) Varsano, N.; Wolf, S. G. Electron Microscopy of Cellular Ultrastructure in Three Dimensions. Current Opinion in Structural Biology 2022, 76, 102444.

(6) Nuñez, J.; Renslow, R.; Cliff, J. B.; Anderton, C. R. NanoSIMS for Biological Applications: Current Practices and Analyses. Biointerphases 2017, 13 (3), 03B301.

(7) De Samber, B.; De Rycke, R.; De Bruyne, M.; Kienhuis, M.; Sandblad, L.; Bohic, S.; Cloetens, P.; Urban, C.; Polerecky, L.; Vincze, L. Effect of Sample Preparation Techniques upon Single Cell Chemical Imaging: A Practical Comparison between Synchrotron Radiation Based X-Ray Fluorescence (SR-XRF) and Nanoscopic Secondary Ion Mass Spectrometry (Nano-SIMS). Anal Chim Acta 2020, 1106, 22–32.

(8) Manisalidis, I.; Stavropoulou, E.; Stavropoulos, A.; Bezirtzoglou, E. Environmental and Health Impacts of Air Pollution: A Review. Front Public Health 2020, 8, 14.

(9) Moreno-Ríos, A. L.; Tejeda-Benítez, L. P.; Bustillo-Lecompte, C. F. Sources, Characteristics, Toxicity, and Control of Ultrafine Particles: An Overview. Geoscience Frontiers 2022, 13 (1), 101147.

(10) Hill, W.; Lim, E. L.; Weeden, C. E.; Lee, C.; Augustine, M.; Chen, K.; Kuan, F.-C.; Marongiu, F.; Evans, E. J.; Moore, D. A.; Rodrigues, F. S.; Pich, O.; Bakker, B.; Cha, H.; Myers, R.; van Maldegem, F.; Boumelha, J.; Veeriah, S.; Rowan, A.; Naceur-Lombardelli, C., et al. Lung Adenocarcinoma Promotion by Air Pollutants. Nature 2023, 616 (7955), 159–167.

(11) You, D. J.; Bonner, J. C. Susceptibility Factors in Chronic Lung Inflammatory Responses to Engineered Nanomaterials. Int J Mol Sci 2020, 21 (19), 7310.

(12) Kempen, P. J.; Kircher, M. F.; de la Zerda, A.; Zavaleta, C. L.; Jokerst, J. V.; Mellinghoff, I. K.; Gambhir, S. S.; Sinclair, R. A Correlative Optical Microscopy and Scanning Electron Microscopy Approach to Locating Nanoparticles in Brain Tumors. Micron 2015, 68, 70–76.

(13) Han, S.; Raabe, M.; Hodgson, L.; Mantell, J.; Verkade, P.; Lasser, T.; Landfester, K.; Weil, T.; Lieberwirth, I. High-Contrast Imaging of Nanodiamonds in Cells by Energy Filtered and Correlative Light-Electron Microscopy: Toward a Quantitative Nanoparticle-Cell Analysis. Nano Lett. 2019, 19 (3), 2178–2185.

(14) Hayashi, Y.; Takamiya, M.; Jensen, P. B.; Ojea-Jiménez, I.; Claude, H.; Antony, C.; Kjaer-Sorensen, K.; Grabher, C.; Boesen, T.; Gilliland, D.; Oxvig, C.; Strähle, U.; Weiss, C. Differential Nanoparticle Sequestration by Macrophages and Scavenger Endothelial Cells Visualized in Vivo in Real-Time and at Ultrastructural Resolution. ACS Nano 2020, 14 (2), 1665–1681.

(15) García-Serradilla, M.; Risco, C. Light and Electron Microscopy Imaging Unveils New Aspects of the Antiviral Capacity of Silver Nanoparticles in Bunyavirus-Infected Cells. Virus Research 2021, 302, 198444.

(16) Kopek, B. G.; Paez-Segala, M. G.; Shtengel, G.; Sochacki, K. A.; Sun, M. G.; Wang, Y.; Xu, C. S.; van Engelenburg, S. B.; Taraska, J. W.; Looger, L. L.; Hess, H. F. Diverse Protocols for Correlative Super-Resolution Fluorescence Imaging and Electron Microscopy of Chemically Fixed Samples. Nat Protoc 2017, 12 (5), 916–946.

(17) Gallagher-Jones, M.; Dias, C. S. B.; Pryor, A.; Bouchmella, K.; Zhao, L.; Lo, Y. H.; Cardoso, M. B.; Shapiro, D.; Rodriguez, J.; Miao, J. Correlative Cellular Ptychography with Functionalized Nanoparticles at the Fe L-Edge. Sci Rep 2017, 7, 4757.

(18) Liu, M.; Li, Q.; Liang, L.; Li, J.; Wang, K.; Li, J.; Lv, M.; Chen, N.; Song, H.; Lee, J.; Shi, J.; Wang, L.; Lal, R.; Fan, C. Real-Time Visualization of Clustering and Intracellular Transport of Gold Nanoparticles by Correlative Imaging. Nat Commun 2017, 8 (1), 15646.

(19) Bhat, S. V.; Sultana, T.; Körnig, A.; McGrath, S.; Shahina, Z.; Dahms, T. E. S. Correlative Atomic Force Microscopy Quantitative Imaging-Laser Scanning Confocal Microscopy Quantifies the Impact of Stressors on Live Cells in Real-Time. Sci Rep 2018, 8 (1), 8305.

(20) Lo Giudice, C.; Yang, J.; Poncin, M. A.; Adumeau, L.; Delguste, M.; Koehler, M.; Evers, K.; Dumitru, A. C.; Dawson, K. A.; Alsteens, D. Nanophysical Mapping of Inflammasome Activation by Nanoparticles via Specific Cell Surface Recognition Events. ACS Nano 2022, 16 (1), 306–316.

(21) Mi, Z.; Chen, C.-B.; Tan, H. Q.; Dou, Y.; Yang, C.; Turaga, S. P.; Ren, M.; Vajandar, S. K.; Yuen, G. H.; Osipowicz, T.; Watt, F.; Bettiol, A. A. Quantifying Nanodiamonds Biodistribution in Whole Cells with Correlative Iono-Nanoscopy. Nat Commun 2021, 12 (1), 4657.

(22) Le Trequesser, Q.; Devès, G.; Saez, G.; Daudin, L.; Barberet, P.; Michelet, C.; Delville, M.-H.; Seznec, H. Single Cell In Situ Detection and Quantification of Metal Oxide Nanoparticles Using Multimodal Correlative Microscopy. Anal. Chem. 2014, 86 (15), 7311–7319.

(23) Podlipec, R.; Punzón-Quijorna, E.; Pirker, L.; Kelemen, M.; Vavpetič, P.; Kavalar, R.; Hlawacek, G.; Štrancar, J.; Pelicon, P.; Fokter, S. K. Revealing Inflammatory Indications Induced by Titanium Alloy Wear Debris in Periprosthetic Tissue by Label-Free Correlative High-Resolution Ion, Electron and Optical Microspectroscopy. Materials 2021, 14 (11), 3048.

(24) Quintana, C.; Wu, T.-D.; Delatour, B.; Dhenain, M.; Guerquin-Kern, J. luc; Croisy, A. Morphological and Chemical Studies of Pathological Human and Mice Brain at the Subcellular Level: Correlation between Light, Electron, and Nanosims Microscopies. Microscopy Research and Technique 2007, 70 (4), 281–295.

(25) Audinot, J.-N.; Georgantzopoulou, A.; Piret, J.-P.; Gutleb, A. C.; Dowsett, D.; Migeon, H. N.; Hoffmann, L. Identification and Localization of Nanoparticles in Tissues by Mass Spectrometry. Surface and Interface Analysis 2013, 45 (1), 230–233.

(26) Vollnhals, F.; Audinot, J.-N.; Wirtz, T.; Mercier-Bonin, M.; Fourquaux, I.; Schroeppel, B.; Kraushaar, U.; Lev-Ram, V.; Ellisman, M. H.; Eswara, S. Correlative Microscopy Combining Secondary Ion Mass Spectrometry and Electron Microscopy: Comparison of Intensity–Hue–Saturation and Laplacian Pyramid Methods for Image Fusion. Anal. Chem. 2017, 89 (20), 10702–10710.

(27) Lovric, J.; Audinot, J.-N.; Wirtz, T. In Situ Correlative Helium Ion Microscopy and Secondary Ion Mass Spectrometry for High-Resolution Nano-Analytics in Life Sciences. Microscopy and Microanalysis 2019, 25 (S2), 1026–1027.

(28) Biesemeier, A.; Castro, O. D.; Serralta, E.; Lovric, J.; Eswara, S.; Audinot, J.-N.; Cambier, S.; Wirtz, T. Correlative Electron Microscopy, High Resolution Ion Imaging and Secondary Ion Mass Spectrometry for High Resolution Nanoanalytics on Biological Tissue. Microscopy and Microanalysis 2020, 26 (S2), 818–820.

(29) De Castro, O.; Biesemeier, A.; Serralta, E.; Bouton, O.; Barrahma, R.; Hoang, Q. H.; Cambier, S.; Taubitz, T.; Klingner, N.; Hlawacek, G.; Pinto, S. D.; Gnauck, P.; Lucas, F.; Bebeacua, C.; Wirtz, T. npSCOPE: A New Multimodal Instrument for In Situ Correlative Analysis of Nanoparticles. Anal. Chem. 2021, 93 (43), 14417–14424.

(30) Le Trequesser, Q.; Saez, G.; Simon, M.; Devès, G.; Daudin, L.; Barberet, P.; Michelet, C.; Delville, M.-H.; Seznec, H. Multimodal Correlative Microscopy for in Situ Detection and Quantification of Chemical Elements in Biological Specimens. Applications to Nanotoxicology. J Chem Biol 2015, 8 (4), 159–167.

(31) James, S. A.; Feltis, B. N.; de Jonge, M. D.; Sridhar, M.; Kimpton, J. A.; Altissimo, M.; Mayo, S.; Zheng, C.; Hastings, A.; Howard, D. L.; Paterson, D. J.; Wright, P. F. A.; Moorhead, G. F.; Turney, T. W.; Fu, J. Quantification of ZnO Nanoparticle Uptake, Distribution, and Dissolution within Individual Human Macrophages. ACS Nano 2013, 7 (12), 10621–10635.

(32) Tardillo Suárez, V.; Gallet, B.; Chevallet, M.; Jouneau, P.-H.; Tucoulou, R.; Veronesi, G.; Deniaud, A. Correlative Transmission Electron Microscopy and High-Resolution Hard X-Ray Fluorescence Microscopy of Cell Sections to Measure Trace Element Concentrations at the Organelle Level. Journal of Structural Biology 2021, 213 (3), 107766.

(33) Malucelli, E.; Iotti, S.; Gianoncelli, A.; Fratini, M.; Merolle, L.; Notargiacomo, A.; Marraccini, C.; Sargenti, A.; Cappadone, C.; Farruggia, G.; Bukreeva, I.; Lombardo, M.; Trombini, C.; Maier, J. A.; Lagomarsino, S. Quantitative Chemical Imaging of the Intracellular Spatial Distribution of Fundamental Elements and Light Metals in Single Cells. Anal Chem 2014, 86 (10), 5108–5115.

(34) Perrin, L.; Carmona, A.; Roudeau, S.; Ortega, R. Evaluation of Sample Preparation Methods for Single Cell Quantitative Elemental Imaging Using Proton or Synchrotron Radiation Focused Beams. J. Anal. At. Spectrom. 2015, 30 (12), 2525–2532.

(35) Jobim, P. F. C.; Iochims dos Santos, C. E.; Dias, J. F.; Kelemen, M.; Pelicon, P.; Mikuš, K. V.; Pascolo, L.; Gianoncelli, A.; Bedolla, D. E.; Rasia-Filho, A. A. Human Neocortex Layer Features Evaluated by PIXE, STIM, and STXM Techniques. Biol Trace Elem Res 2023, 201 (2), 592–602.

(36) Pongrac, P.; Vogel-Mikuš, K.; Jeromel, L.; Vavpetič, P.; Pelicon, P.; Kaulich, B.; Gianoncelli, A.; Eichert, D.; Regvar, M.; Kreft, I. Spatially Resolved Distributions of the Mineral Elements in the Grain of Tartary Buckwheat (*Fagopyrum Tataricum*). Food Research International 2013, 54 (1), 125–131.

(37) Bischof, J.; Fletcher, G.; Verkade, P.; Kuntner, C.; Fernandez-Rodriguez, J.; Chaabane, L.; Rose, L. A.; Walter, A.; Vandenbosch, M.; van Zandvoort, M. A. M. J.; Zaritsky, A.; Keppler, A.; Parsons, M. Multimodal Bioimaging across Disciplines and Scales: Challenges, Opportunities and Breaking down Barriers. npj Imaging 2024, 2 (1), 1–6.

(38) Oberdörster, G.; Maynard, A.; Donaldson, K.; Castranova, V.; Fitzpatrick, J.; Ausman, K.; Carter, J.; Karn, B.; Kreyling, W.; Lai, D.; Olin, S.; Monteiro-Riviere, N.; Warheit, D.; Yang, H.; A report from the ILSI Research Foundation/Risk Science Institute Nanomaterial Toxicity Screening Working Group. Principles for Characterizing the Potential Human Health Effects from Exposure to Nanomaterials: Elements of a Screening Strategy. Particle and Fibre Toxicology 2005, 2 (1), 8.

(39) Kokot, H.; Kokot, B.; Sebastijanović, A.; Voss, C.; Podlipec, R.; Zawilska, P.; Berthing, T.; Ballester-López, C.; Danielsen, P. H.; Contini, C.; Ivanov, M.; Krišelj, A.; Čotar, P.; Zhou, Q.; Ponti, J.; Zhernovkov, V.; Schneemilch, M.; Doumandji, Z.; Pušnik, M.; Umek, P.; Pajk, S., et al., J. Prediction of Chronic Inflammation for Inhaled Particles: The Impact of Material Cycling and Quarantining in the Lung Epithelium. Advanced Materials 2020, 32 (47), 2003913.

(40) Danielsen, P. H.; Knudsen, K. B.; Štrancar, J.; Umek, P.; Koklič, T.; Garvas, M.; Vanhala, E.; Savukoski, S.; Ding, Y.; Madsen, A. M.; Jacobsen, N. R.; Weydahl, I. K.; Berthing, T.; Poulsen, S. S.; Schmid, O.; Wolff, H.; Vogel, U. Effects of Physicochemical Properties of TiO2 Nanomaterials for Pulmonary Inflammation, Acute Phase Response and Alveolar Proteinosis in Intratracheally Exposed Mice. Toxicology and Applied Pharmacology 2020, 386, 114830.

(41) Podlipec, R.; Sebastijanovič, A.; Cassidy, H.; Han, L.; Danielsen, P.; Čotar, P.; Kokot, H.; Koser, B. K.; Vencelj, A.; Pirker, L.; Umek, P.; Hlawacek, G.; Heller, R.; Matallanas, D.; Pelicon, P.; Stoeger, T.; Urbančič, I.; Vogel, U.; Koklič, T.; Štrancar, J. Binucleated Cell Formation and Oncogene Expression after Particulate Matter Exposure Is Preceded by Microtubule Disruption, Dysregulated Cell Cycle, Prolonged Mitosis, and Septin Binding. bioRxiv, 10.1101/2024.01.07.574515.

(42) Zhang, Y.; Huang, T.; Jorgens, D. M.; Nickerson, A.; Lin, L.-J.; Pelz, J.; Gray, J. W.; López, C. S.; Nan, X. Quantitating Morphological Changes in Biological Samples during Scanning Electron Microscopy Sample Preparation with Correlative Super-Resolution Microscopy. PLoS One 2017, 12 (5), e0176839.

(43) Li, Y.; Almassalha, L. M.; Chandler, J. E.; Zhou, X.; Stypula-Cyrus, Y. E.; Hujsak, K. A.; Roth, E. W.; Bleher, R.; Subramanian, H.; Szleifer, I.; Dravid, V. P.; Backman, V. The Effects of Chemical Fixation on the Cellular Nanostructure. Exp Cell Res 2017, 358 (2), 253–259.

(44) Murk, J. L. a. N.; Posthuma, G.; Koster, A. J.; Geuze, H. J.; Verkleij, A. J.; Kleijmeer, M. J.; Humbel, B. M. Influence of Aldehyde Fixation on the Morphology of Endosomes and Lysosomes: Quantitative Analysis and Electron Tomography. Journal of Microscopy 2003, 212 (1), 81–90.

(45) Ebersold, H. R.; Cordier, J. L.; Lüthy, P. Bacterial Mesosomes: Method Dependent Artifacts. Arch Microbiol 1981, 130 (1), 19–22.

(46) Idziak, A.; Inavalli, V. V. G. K.; Bancelin, S.; Arizono, M.; Nägerl, U. V. The Impact of Chemical Fixation on the Microanatomy of Mouse Organotypic Hippocampal Slices. eNeuro 2023, 10 (9).

(47) Parlanti, P.; Cappello, V. Microscopes, Tools, Probes, and Protocols: A Guide in the Route of Correlative Microscopy for Biomedical Investigation. Micron 2022, 152, 103182.

(48) Kokot, B.; Kokot, H.; Umek, P.; van Midden, K. P.; Pajk, S.; Garvas, M.; Eggeling, C.; Koklič, T.; Urbančič, I.; Štrancar, J. How to Control Fluorescent Labeling of Metal Oxide Nanoparticles for Artefact-Free Live Cell Microscopy. Nanotoxicology 2021, 15 (8), 1102–1123.

44. Suhling, K. et al. Fluorescence Lifetime Imaging. in Handbook of Photonics for Biomedical Engineering (eds. Ho, A. H.-P., Kim, D. & Somekh, M. G.) 1–50 (Springer Netherlands, Dordrecht, 2021).

(50) Strickfaden, H. Reflections on the Organization and the Physical State of Chromatin in Eukaryotic Cells. Genome 2021, 64 (4), 311–325.

(51) Shrestha, D.; Jenei, A.; Nagy, P.; Vereb, G.; Szöllősi, J. Understanding FRET as a Research Tool for Cellular Studies. Int J Mol Sci 2015, 16 (4), 6718–6756.

(52) Miller, K. N.; Victorelli, S. G.; Salmonowicz, H.; Dasgupta, N.; Liu, T.; Passos, J. F.; Adams, P. D. Cytoplasmic DNA: Sources, Sensing, and Role in Aging and Disease. Cell 2021, 184 (22), 5506–5526.

(53) Pol, E. van der; Böing, A. N.; Harrison, P.; Sturk, A.; Nieuwland, R. Classification, Functions, and Clinical Relevance of Extracellular Vesicles. Pharmacol Rev 2012, 64 (3), 676–705.

(54) Goldstein, J. I.; Newbury, D. E.; Michael, J. R.; Ritchie, N. W. M.; Scott, J. H. J.; Joy, D. C. Energy Dispersive X-Ray Spectrometry: Physical Principles and User-Selected Parameters. In Scanning Electron Microscopy and X-Ray Microanalysis; Goldstein, J. I., Newbury, D. E., Michael, J. R., Ritchie, N. W. M., Scott, J. H. J., Joy, D. C., Eds.; Springer: New York, NY, 2018; pp 209–234.

(55) Urbančič, I.; Garvas, M.; Kokot, B.; Majaron, H.; Umek, P.; Cassidy, H.; Škarabot, M.; Schneider, F.; Galiani, S.; Arsov, Z.; Koklic, T.; Matallanas, D.; Čeh, M.; Muševič, I.; Eggeling, C.; Štrancar, J. Nanoparticles Can Wrap Epithelial Cell Membranes and Relocate Them Across the Epithelial Cell Layer. Nano Lett. 2018, 18 (8), 5294–5305.

(56) Elzanowska, J.; Semira, C.; Costa-Silva, B. DNA in Extracellular Vesicles: Biological and Clinical Aspects. Molecular Oncology 2020, 15 (6), 1701.

(57) Wyllie, A. H.; Kerr, J. F.; Currie, A. R. Cell Death: The Significance of Apoptosis. Int Rev Cytol 1980, 68, 251–306.

(58) Karbowski, M.; Youle, J. R., Dynamics of mitochondrial morphology in healthy cells and during apoptosis, Cell Death Differ. 2003, 10, 870–880.

(59) Zhang, X.; Wang, F.; Liu, B.; Kelly, E. Y.; Servos, M. R.; Liu, J. Adsorption of DNA Oligonucleotides by Titanium Dioxide Nanoparticles. Langmuir 2014, 30 (3), 839–845.

(60) Bischoff, M.; Biriukov, D.; Předota, M.; Roke, S.; Marchioro, A. Surface Potential and Interfacial Water Order at the Amorphous TiO2 Nanoparticle/Aqueous Interface. J. Phys. Chem. C 2020, 124 (20), 10961–10974.

(61) Shukla, R. K.; Badiye, A.; Vajpayee, K.; Kapoor, N. Genotoxic Potential of Nanoparticles: Structural and Functional Modifications in DNA. Front Genet 2021, 12, 728250.

(62) Morrow, P. Possible Mechanisms to Explain Dust Overloading of the Lungs. Fundamental and Applied Toxicology 1988, 10 (3), 369–384.

(63) Sachet, M.; Liang, Y. Y.; Oehler, R. The Immune Response to Secondary Necrotic Cells. Apoptosis 2017, 22 (10), 1189–1204.

(64) Degen, M.; Santos, J. C.; Pluhackova, K.; Cebrero, G.; Ramos, S.; Jankevicius, G.; Hartenian, E.; Guillerm, U.; Mari, S. A.; Kohl, B.; Müller, D. J.; Schanda, P.; Maier, T.; Perez, C.; Sieben, C.; Broz, P.; Hiller, S. Structural Basis of NINJ1-Mediated Plasma Membrane Rupture in Cell Death. Nature 2023, 618 (7967), 1065–1071.

(65) Zhao, C.; Pu, W.; Niu, M.; Wazir, J.; Song, S.; Wei, L.; Li, L.; Su, Z.; Wang, H. Respiratory Exposure to PM2.5 Soluble Extract Induced Chronic Lung Injury by Disturbing the Phagocytosis Function of Macrophage. Environ Sci Pollut Res 2022, 29 (10), 13983–13997.

(66) Gustafson, H. H.; Holt-Casper, D.; Grainger, D. W.; Ghandehari, H. Nanoparticle Uptake: The Phagocyte Problem. Nano Today 2015, 10 (4), 487–510.

(67) Chen, Q.; Wang, N.; Zhu, M.; Lu, J.; Zhong, H.; Xue, X.; Guo, S.; Li, M.; Wei, X.; Tao, Y.; Yin, H. TiO2 Nanoparticles Cause Mitochondrial Dysfunction, Activate Inflammatory Responses, and Attenuate Phagocytosis in Macrophages: A Proteomic and Metabolomic Insight. Redox Biology 2018, 15, 266–276.

(68) Gianoncelli, A.; Bonanni, V.; Gariani, G.; Guzzi, F.; Pascolo, L.; Borghes, R.; Billè, F.; Kourousias, G. Soft X-Ray Microscopy Techniques for Medical and Biological Imaging at TwinMic—Elettra. Applied Sciences 2021, 11 (16), 7216.

(69) Urbančič, I.; Arsov, Z.; Ljubetič, A.; Biglino, D.; Strancar, J. Bleaching-Corrected Fluorescence Microspectroscopy with Nanometer Peak Position Resolution. Opt Express 2013, 21 (21), 25291–25306.

53. Solvent and Environmental Effects. in Principles of Fluorescence Spectroscopy (ed. Lakowicz, J. R.) 205–235 (Springer US, Boston, MA, 2006).

(71) Berman, E. S. F.; Fortson, S. L.; Checchi, K. D.; Wu, L.; Felton, J. S.; Wu, K. J. J.; Kulp, K. S. Preparation of Single Cells for Imaging/Profiling Mass Spectrometry. Journal of the American Society for Mass Spectrometry 2008, 19 (8), 1230–1236.

(72) Liu, Y.; Tan, J.; Thomas, A.; Ou-Yang, D.; Muzykantov, V. R. The Shape of Things to Come: Importance of Design in Nanotechnology for Drug Delivery. Ther Deliv 2012, 3 (2), 181–194.

(73) Lesniak, A.; Fenaroli, F.; Monopoli, M. P.; Åberg, C.; Dawson, K. A.; Salvati, A. Effects of the Presence or Absence of a Protein Corona on Silica Nanoparticle Uptake and Impact on Cells. ACS Nano 2012, 6 (7), 5845–5857.

(74) First-Principles Study of Na Insertion at TiO2 Anatase Surfaces: New Hints for Na-Ion Battery Design. Nanoscale Advances 2020, 2 (7), 2745–2751.

(75) Shriver, Z.; Capila, I.; Venkataraman, G.; Sasisekharan, R. Heparin and Heparan Sulfate: Analyzing Structure and Microheterogeneity. Handb Exp Pharmacol 2012, No. 207, 159– 176.

(76) Farrugia, B. L.; Lord, M. S.; Melrose, J.; Whitelock, J. M. The Role of Heparan Sulfate in Inflammation, and the Development of Biomimetics as Anti-Inflammatory Strategies. J Histochem Cytochem 2018, 66 (4), 321–336.

(77) Lu, M.; Kaminski, C. F.; Schierle, G. S. K. Advanced Fluorescence Imaging of in Situ Protein Aggregation. Phys Biol 2020, 17 (2), 021001.

(78) Rustom, A.; Saffrich, R.; Markovic, I.; Walther, P.; Gerdes, H.-H. Nanotubular Highways for Intercellular Organelle Transport. Science 2004, 303 (5660), 1007–1010.

(79) Donovan, A.; Lima, C. A.; Pinkus, J. L.; Pinkus, G. S.; Zon, L. I.; Robine, S.; Andrews, N. C. The Iron Exporter Ferroportin/Slc40a1 Is Essential for Iron Homeostasis. Cell Metabolism 2005, 1 (3), 191–200.

(80) Nakamura, T.; Naguro, I.; Ichijo, H. Iron Homeostasis and Iron-Regulated ROS in Cell Death, Senescence and Human Diseases. Biochimica et Biophysica Acta (BBA) - General Subjects 2019, 1863 (9), 1398–1409.

(81) Nano, G. V.; Strathmann, T. J. Ferrous Iron Sorption by Hydrous Metal Oxides. Journal of Colloid and Interface Science 2006, 297 (2), 443–454.

(82) Mayer, K. M.; Hafner, J. H. Localized Surface Plasmon Resonance Sensors. Chem. Rev. 2011, 111 (6), 3828–3857.

(83) Tang, L.; Casas, J.; Venkataramasubramani, M. Magnetic Nanoparticle Mediated Enhancement of Localized Surface Plasmon Resonance for Ultrasensitive Bioanalytical Assay in Human Blood Plasma. Anal Chem 2013, 85 (3), 1431–1439.

(84) Li, J.; Krasavin, A. V.; Webster, L.; Segovia, P.; Zayats, A. V.; Richards, D. Spectral Variation of Fluorescence Lifetime near Single Metal Nanoparticles. Sci Rep 2016, 6 (1), 21349.

(85) Li, J.; Cao, F.; Yin, H.; Huang, Z.; Lin, Z.; Mao, N.; Sun, B.; Wang, G. Ferroptosis: Past, Present and Future. Cell Death Dis 2020, 11 (2), 1–13.

(86) Danylchuk, D. I.; Jouard, P.-H.; Klymchenko, A. S. Targeted Solvatochromic Fluorescent Probes for Imaging Lipid Order in Organelles under Oxidative and Mechanical Stress. J Am Chem Soc 2021, 143 (2), 912–924.

(87) Ryan, E.; Mockros, L.; Weisel, J.; Lorand, L. Structural Origins of Fibrin Clot Rheology. Biophys J 1999, 77 (5), 2813–2826.

(88) Li, W.; Sigley, J.; Pieters, M.; Helms, C. C.; Nagaswami, C.; Weisel, J. W.; Guthold, M. Fibrin Fiber Stiffness Is Strongly Affected by Fiber Diameter, but Not by Fibrinogen Glycation. Biophys J 2016, 110 (6), 1400–1410.

(89) Idell, S.; James, K. K.; Levin, E. G.; Schwartz, B. S.; Manchanda, N.; Maunder, R. J.; Martin, T. R.; McLarty, J.; Fair, D. S. Local Abnormalities in Coagulation and Fibrinolytic Pathways Predispose to Alveolar Fibrin Deposition in the Adult Respiratory Distress Syndrome. J Clin Invest 1989, 84 (2), 695–705.

(90) Idell, S. Coagulation, Fibrinolysis, and Fibrin Deposition in Acute Lung Injury. Crit Care Med 2003, 31 (4 Suppl), S213–220.

(91) Guadiz, G.; Sporn, L. A.; Goss, R. A.; Lawrence, S. O.; Marder, V. J.; Simpson-Haidaris, P. J. Polarized Secretion of Fibrinogen by Lung Epithelial Cells. Am J Respir Cell Mol Biol 1997, 17 (1), 60–69.

(92) Bastarache, J. A.; Sebag, S. C.; Grove, B. S.; Ware, L. B. IFN-γ and TNF-α Act Synergistically to Upregulate Tissue Factor in Alveolar Epithelial Cells. Exp Lung Res 2011, 37 (8), 509–517.

(93) Naji, A.; Muzembo, B. A.; Yagyu, K.; Baba, N.; Deschaseaux, F.; Sensebé, L.; Suganuma, N. Endocytosis of Indium-Tin-Oxide Nanoparticles by Macrophages Provokes Pyroptosis Requiring NLRP3-ASC-Caspase1 Axis That Can Be Prevented by Mesenchymal Stem Cells. Sci Rep 2016, 6 (1), 26162.

(94) Erdem, J. S.; Závodná, T.; Ervik, T. K.; Skare, Ø.; Hron, T.; Anmarkrud, K. H.; Kuśnierczyk, A.; Catalán, J.; Ellingsen, D. G.; Topinka, J.; Zienolddiny-Narui, S. High Aspect Ratio Nanomaterial-Induced Macrophage Polarization Is Mediated by Changes in miRNA Levels. Front Immunol 2023, 14, 1111123.

(95) Garvas, M.; Testen, A.; Umek, P.; Gloter, A.; Koklic, T.; Strancar, J. Protein Corona Prevents TiO2 Phototoxicity. PLoS One 2015, 10 (6), e0129577.

(96) Baranowska-Wójcik, E.; Szwajgier, D.; Oleszczuk, P.; Winiarska-Mieczan, A. Effects of Titanium Dioxide Nanoparticles Exposure on Human Health—a Review. Biol Trace Elem Res 2020, 193 (1), 118–129.

(97) Gutierrez, C. T.; Loizides, C.; Hafez, I.; Brostrøm, A.; Wolff, H.; Szarek, J.; Berthing, T.; Mortensen, A.; Jensen, K. A.; Roursgaard, M.; Saber, A. T.; Møller, P.; Biskos, G.; Vogel, U. Acute Phase Response Following Pulmonary Exposure to Soluble and Insoluble Metal Oxide Nanomaterials in Mice. Particle and Fibre Toxicology 2023, 20 (1), 4.

(98) Wang, J. J.; Sanderson, B. J. S.; Wang, H. Cyto- and Genotoxicity of Ultrafine TiO2 Particles in Cultured Human Lymphoblastoid Cells. Mutation Research/Genetic Toxicology and Environmental Mutagenesis 2007, 628 (2), 99–106.

(99) Schmid, O.; Stoeger, T. Surface Area Is the Biologically Most Effective Dose Metric for Acute Nanoparticle Toxicity in the Lung. Journal of Aerosol Science 2016, 99, 133–143.

(100) Jackson, P.; Halappanavar, S.; Hougaard, K. S.; Williams, A.; Madsen, A. M.; Lamson, J. S.; Andersen, O.; Yauk, C.; Wallin, H.; Vogel, U. Maternal Inhalation of Surface-Coated Nanosized Titanium Dioxide (UV-Titan) in C57BL/6 Mice: Effects in Prenatally Exposed Offspring on Hepatic DNA Damage and Gene Expression. Nanotoxicology 2013, 7 (1), 85–96.

(101) Tivol, W. F.; Briegel, A.; Jensen, G. J. An Improved Cryogen for Plunge Freezing. Microsc Microanal 2008, 14 (5), 375–379.

(102) Becker, W. The Bh TCSPC Handbook 10th Edition, 2023.

(103) Podlipec, R.; Mur, J.; Petelin, J.; Štrancar, J.; Petkovšek, R. Two-Photon Retinal Theranostics by Adaptive Compact Laser Source. Appl. Phys. A 2020, 126 (6), 405.

(104) Podlipec, R.; Mur, J.; Petelin, J.; Štrancar, J.; Petkovšek, R. Method for Controlled Tissue Theranostics Using a Single Tunable Laser Source. Biomed Opt Express 2021, 12 (9), 5881–5893.

(105) Helium Ion Microscopy; Hlawacek, G., Gölzhäuser, A., Eds.; NanoScience and Technology; Springer International Publishing: Cham, 2016.

(106) Höflich, K.; Hobler, G.; Allen, F. I.; Wirtz, T.; Rius, G.; McElwee-White, L.; Krasheninnikov, A. V.; Schmidt, M.; Utke, I.; Klingner, N.; Osenberg, M.; Córdoba, R.; Djurabekova, F.; Manke, I.; Moll, P.; Manoccio, M.; De Teresa, J. M.; Bischoff, L.; Michler, J.; De Castro, O., et al. Roadmap for Focused Ion Beam Technologies. Applied Physics Reviews 2023, 10 (4), 041311.

(107) Hlawacek, G.; Veligura, V.; van Gastel, R.; Poelsema, B. Helium Ion Microscopy. Journal of Vacuum Science & Technology B 2014, 32 (2), 020801. 10.1116/1.4863676.

(108) Schmidt, M.; Byrne, J. M.; Maasilta, I. J. Bio-Imaging with the Helium-Ion Microscope: A Review. Beilstein J. Nanotechnol. 2021, 12 (1), 1–23.

(109) Gianoncelli, A.; Kourousias, G.; Merolle, L.; Altissimo, M.; Bianco, A. Current Status of the TwinMic Beamline at Elettra: A Soft X-Ray Transmission and Emission Microscopy Station. J Synchrotron Radiat 2016, 23 (Pt 6), 1526–1537.

(110) Gianoncelli, A.; Morrison, G. R.; Kaulich, B.; Bacescu, D.; Kovac, J. Scanning Transmission X-Ray Microscopy with a Configurable Detector. Applied Physics Letters 2006, 89 (25), 251117.

(111) Gianoncelli, A.; Kourousias, G.; Stolfa, A.; Kaulich, B. Recent Developments at the TwinMic Beamline at ELETTRA: An 8 SDD Detector Setup for Low Energy X-Ray Fluorescence. J. Phys.: Conf. Ser. 2013, 425 (18), 182001.

(112) Solé, V. A.; Papillon, E.; Cotte, M.; Walter, Ph.; Susini, J. A Multiplatform Code for the Analysis of Energy-Dispersive X-Ray Fluorescence Spectra. Spectrochimica Acta Part B: Atomic Spectroscopy 2007, 62 (1), 63–68.

