## Supporting Information for "Novel correlative microscopy approach for nano-bio interface studies of nanoparticle-induced lung epithelial cell damage"

**This file includes:**

Supplementary Figures 1 - 8

Supplementary Comments #1 - #2

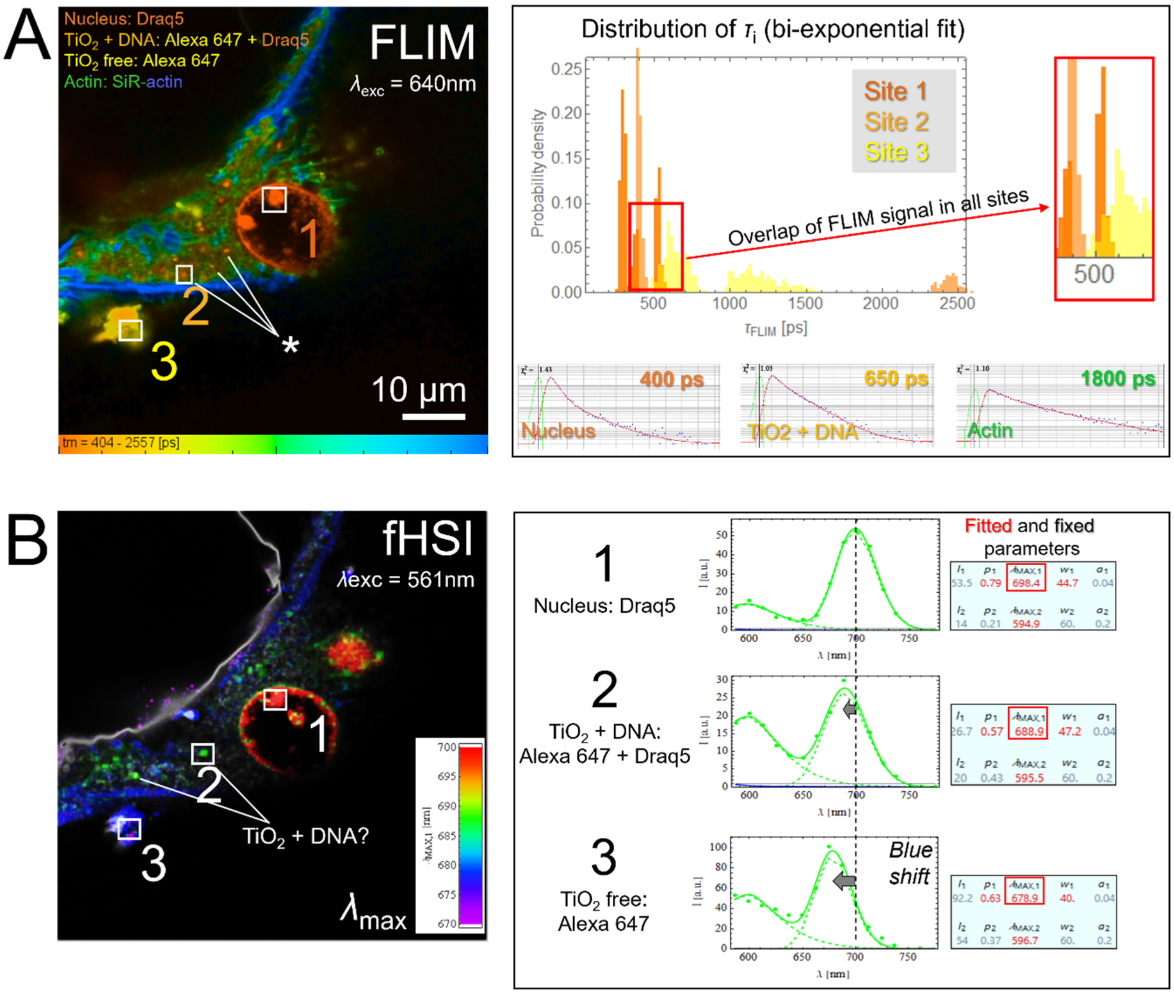


**Figure S1.** Combined fluorescence lifetime imaging microscopy (FLIM) (A) and fluorescence hyperspectral imaging (fHSI) (B) of lung epithelial cells exposed to $TiO_2$ NTs confirms a local and widespread distribution of DNA in the cytoplasm, apparently co-localized with the nanoparticles (marked with an asterisk). This is supported by (1) the partial overlap of the $\tau_i$ distribution for all the measured sites (1-3) at around 500 ps (A), characteristic for nuclear/DNA dye, and (2) by the distinct spectral shifts of the second fitted component, where the one at site 2 ($\lambda_{max}$ = 688 nm) is most likely a superposition of the spectral components at site 1 ($\lambda_{max}$ = 698 nm, characteristic for DNA/Draq5) and site 3 (($\lambda_{max}$ = 678 nm, characteristic for TiO2/Alexa647).

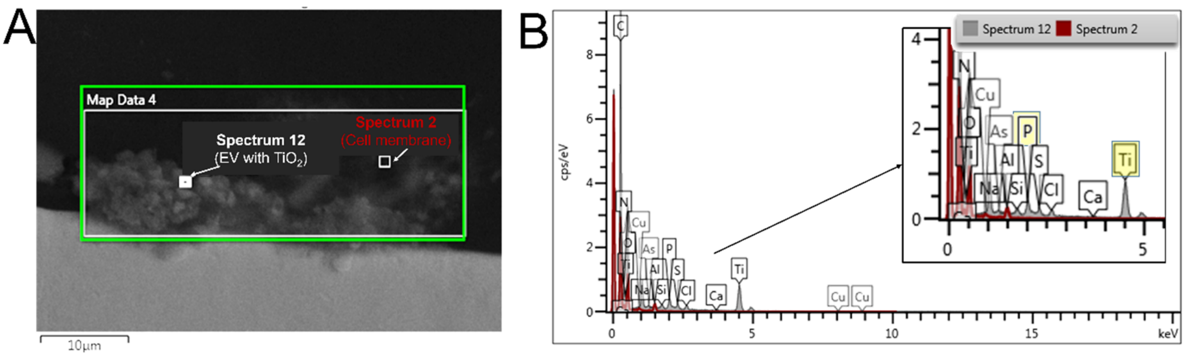


**Figure S2.** An example of SEM imaging (A) and the following spectral analysis of the elemental distribution within the marked spots, one inside the large EV with TiO2 and in the control cell membrane (B).

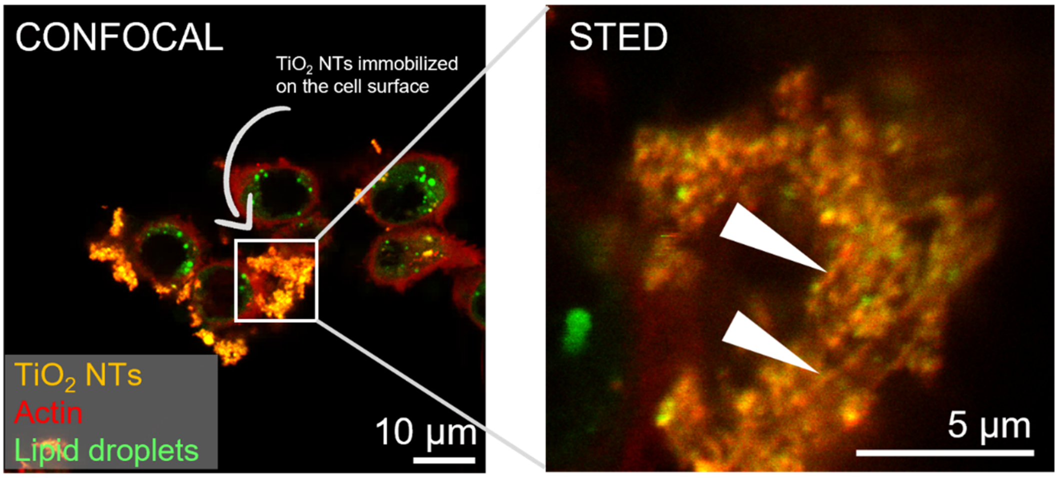


**Figure S3**. High-resolution confocal and super-resolution STED microscopy performed on TiO2-bio composites immobilized on the cellular surface of lung epithelial LA4 cells. STED microscopy in a thin, submicron, image plane on the cellular surface uncovers the heterogeneous structure with the actin fibers intertwined and spread across the TiO2-rich composite (white arrows).

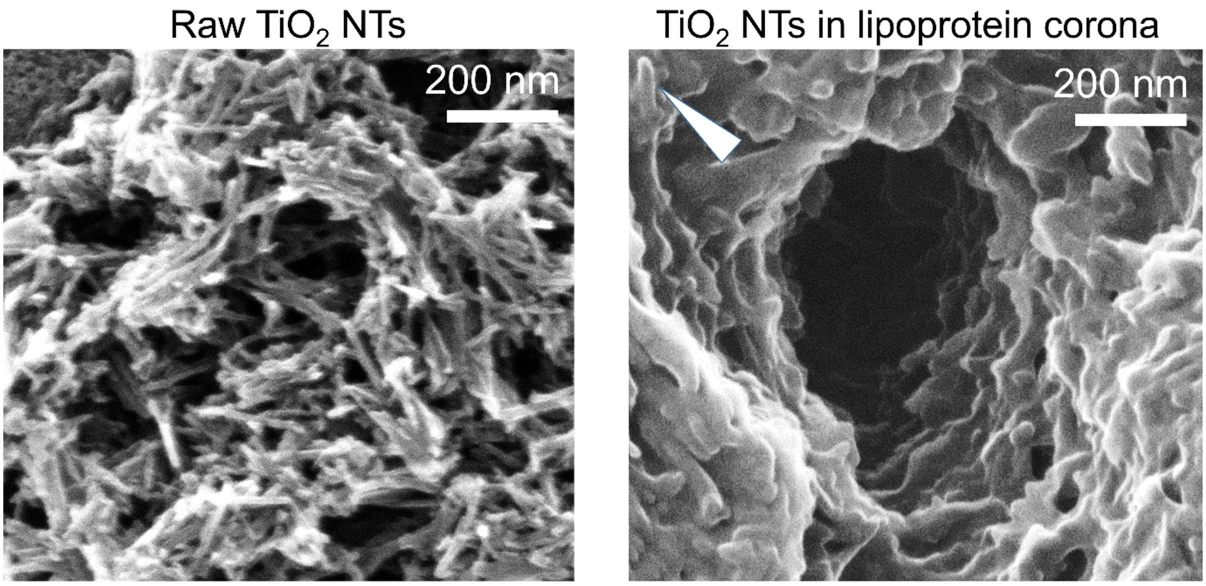


**Figure S4**. Clear demonstration of the extensive biomolecule binding to the immobilized $TiO_2$ NTs on the lung epithelial cell surface, forming the so-called lipoprotein corona (right image) with a few nm layer (see the arrow). In contrast, uncoated $TiO_2$ NTs are shown in the left image as a control. Precise quantification of the biological coating was achieved using a sub-nanometer resolution helium ion microscopy (HIM), which, unlike SEM, does not require a conductive coating on these highly insulating samples—thus preserving their original surface properties.

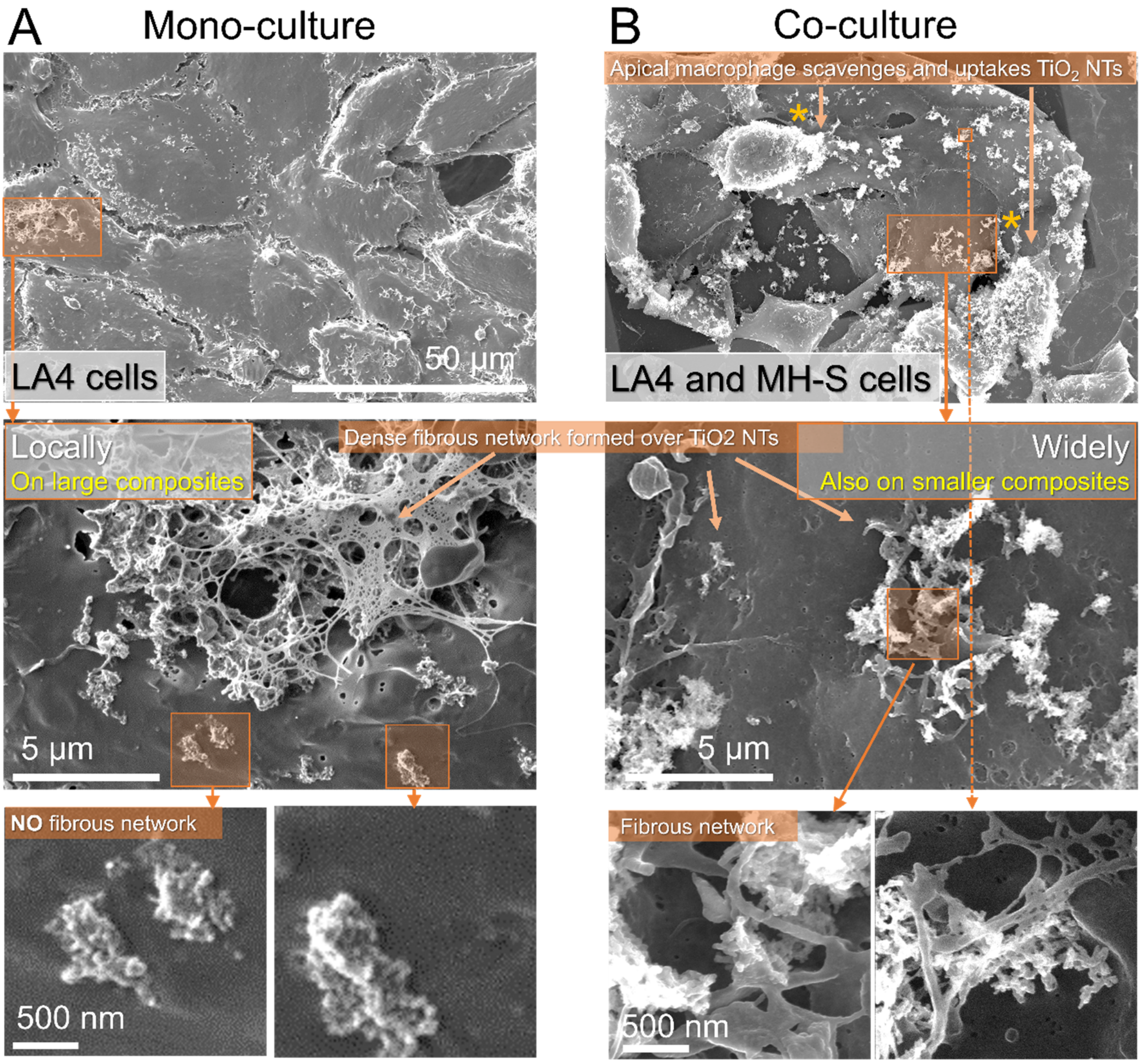


**Figure S5**. Fibrous structure formation localized on the TiO2-bio composites immobilized on the lung epithelial surface, as one of the potential crucial initiating modes of action in inflammatory cell response, measured in mono-culture (A) and co-culture with innate immune cells (B). A) Formation of a dense fibrous network spread locally over the surface of larger TiO2-bio composites, sparring the smaller ones (bottom images). B) Formation of a dense fibrous network spread over the surface basically all TiO2-bio composites (bottom images) which might be due to the presence of an activated MH-S macrophages, which release cytokines, such as TNFα and may induce or boost the formation of fibrin fibers. Apical macrophages are marked with an asterisk and shown magnified in the Fig. S5. Imaging was performed on FEI Helios Nanolab SEM.

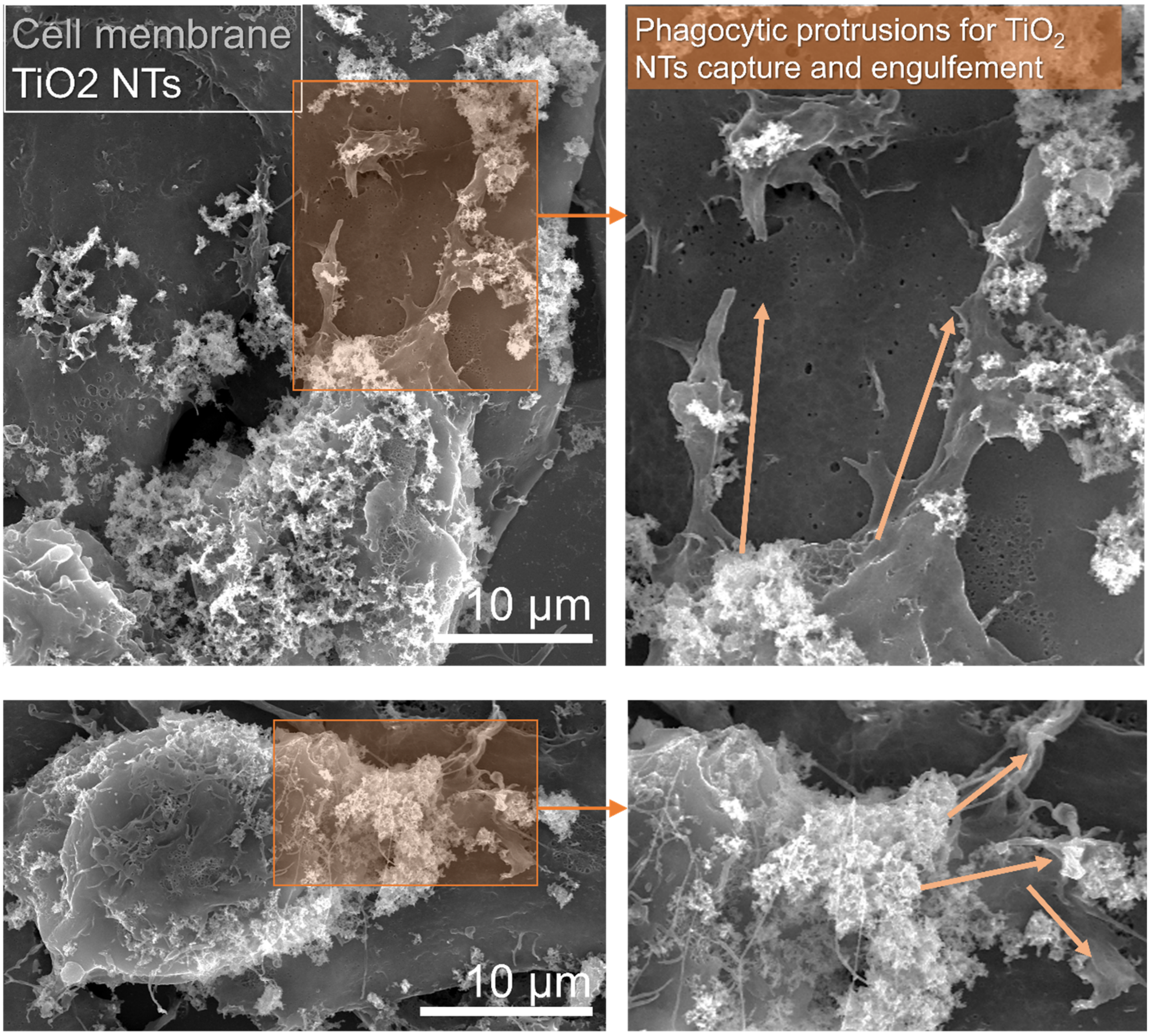


**Figure S6.** Apical macrophages on the top of lung epithelial LA4 cells with the extended phagocytic protrusions (marked with the arrows) capturing and internalizing $TiO_2$ NTs. Nanoparticles or nano-bio composites are shown with white contrast due to high secondary electrons (SE) yield compared to organic matter using FEI Helios Nanolab SEM.

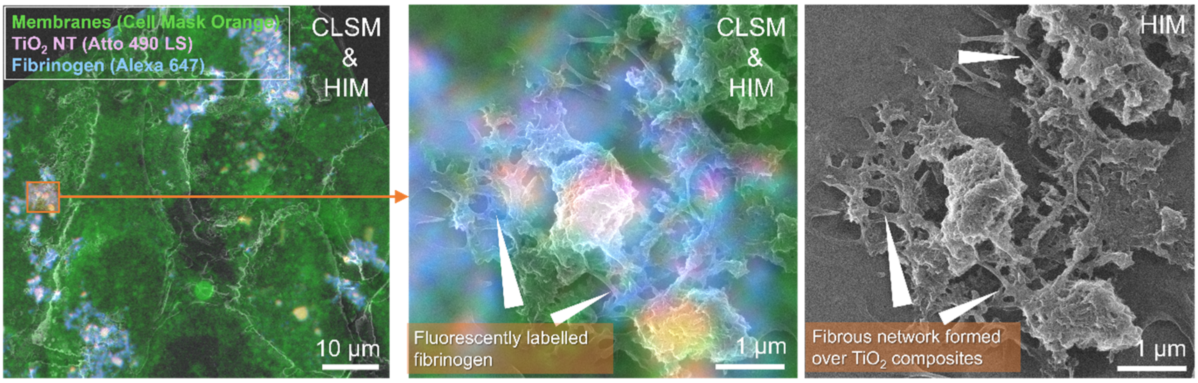


**Figure S7.** Measuring fibrinogen co-localization with the fibrous structures formed locally over $TiO_2$-bio composites with the correlated 3-channel CLSM and ultra-high resolution HIM. Despite not perfect co-localization due to two orders of magnitude poorer resolution of CLSM and slight sample displacement during rapid cryo-fixation and freeze drying, the results still indicate on fibrin presence in the fibrous structures.

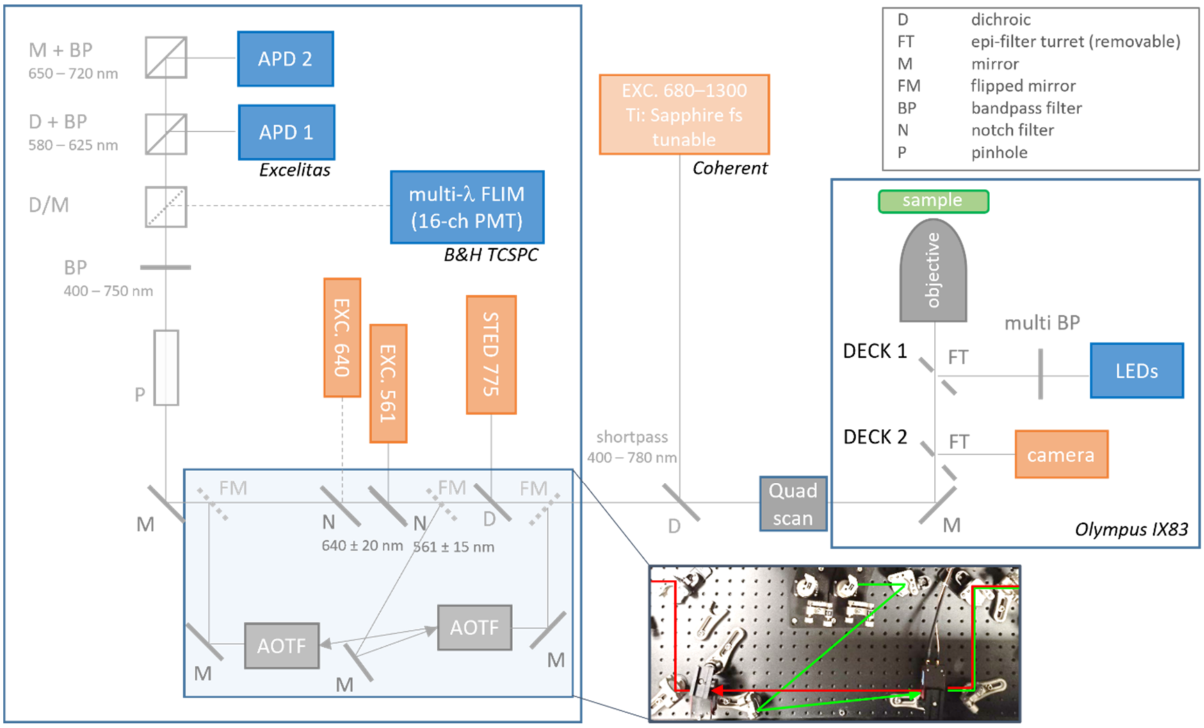


**Figure S8**. Scheme of the custom-built optical setup for confocal laser scanning microscopy (CLSM), super-resolution microscopy (STED), fluorescence lifetime imaging microscopy (FLIM) and fluorescence hyperspectral imaging (fHSI). Light sources, pulsed diode lasers (561nm and 640nm) and LEDs are shown in orange color, detectors (avalanche photodiodes – APDs and 16-channel photomultiplier tubes - PMTs) in blue. In order to avoid partial blocking of photons on the notch filters in our de-scanned detection setup, which distorts the acquired fluorescence spectra, an optical bypass for collected photons was developed using a set of acousto-optical tunable filters (AOTF, Opto-electronic).

### Appendix

#### Supplementary Comment #1

The $TiO_2$ particles are worldwide and one of the most ubiquitous industrial NMs, widely used in paints, plastics, cosmetics, electronics, etc. Their widespread use can lead to locally high concentrations of dust in the air, where they remain suspended for extended periods due to their small size, increasing human exposure and thus undesirable health effects. They have also been classified as Group 2b carcinogens by the International Agency for Research on Cancer (IARC). To this end, the National Institute for Occupational Safety and Health (NIOSH) has recommended airborne exposure limits of 2.4 mg/m$^3$ for fine $TiO_2$ and 0.3 mg/m$^3$ for ultrafine (including engineered nanoscale) $TiO_2$, as time-weighted average (TWA) concentrations for up to 10 hr/day during a 40-hour work week.[1] These recommendations represent levels that are estimated to reduce the risk of lung cancer to less than 1 in 1,000 over a working lifetime. The recommendations are based on the use of chronic inhalation studies in rats to predict lung tumor risks in humans.

The concentration of $TiO_2$ NTs used in our study was 10 µg/ml, resulting in a total surface dose of 3 µg/cm$^2$. To put this into perspective, when considering the average alveolar surface area of an adult human of 80 m$^2$, 2.4 g of $TiO_2$ would have to be deposited to achieve a surface dose of 3 µg/cm$^2$. Based on an average tidal volume of air per breathing cycle (500 mL) and the NIOSH occupational exposure safety limits for ultrafine $TiO_2$, the dose we used is equivalent to 0.3 mg/m$^3$ exposure over a 10-year working days period, based on the 8-hour time-weighted average occupational exposure. In the case of highly probable partial retention, translocation and clearance of nanoparticles, the used dose is estimated to accumulate over a working lifetime at defined airborne exposure limits.

#### Supplementary Comment #2

Future prospects in correlative microscopy workflow point to improved automation, starting with reducing the role of the operator to limit his experimental variability in sample handling and further sample processing. The latter has recently been addressed with several semi-automated registration and image alignment software supports[2–4], but further integration with hardware development is required. However, this transition is significantly hampered by the high cost of the integrated systems, which on the other hand significantly reduce experimental variability by automated sample transfer between stages, such as in several cryo-CLEM systems from Leica Microsystems or Zeiss. An alternative strategy to these state-of-the-art integrated systems, where the extremely high costs typically outweigh the benefits, is to improve the automation of existing workflows, which can be much more cost effective.

One approach is the development of high-throughput imaging, which involves automated, trained detection, tracking and subsequent analysis of the fluorescently labelled cellular features of interest on live cells using various free and open-source software tools such as CellProfiler[5] or Cellpose.[6] Easily stored coordinates and positions of the studied features/structures can then be used as input data for stage translation in other instruments (e.g. electron microscope) after successful registration of the key fiducial point in the customized sample holders. Measurements over the multiple sites, when not too time consuming as may be in the case of a large field-of-view scan in SR XRF, can provide sufficient statistical data for meaningful interpretation and insight into the biological system being studied. In principle, a high-throughput approach can also partially solve the problem of very likely incomplete sample preservation due to potential structural changes and sample deformations during the various steps from chemical fixation, cryo-fixation to final embedding and sectioning. Multiple data sets can be very useful to identify different patterns of compromised samples and to remove unwanted data in the analysis to improve overall data reliability. However, there is a need to better optimize the control of all physical variables throughout sample preparation, which currently relies heavily on operator accuracy in sample handling during blotting and plating. This can be addressed by partial automation, such as the integrated approach used in the new Linkam Plunger.[7]

### References


(1) *Current Intelligence Bulletin 63: Occupational Exposure to Titanium Dioxide.*; U.S. Department of Health and Human Services, Public Health Service, Centers for Disease Control and Prevention, National Institute for Occupational Safety and Health, 2011.

(2) Paul-Gilloteaux, P.; Heiligenstein, X.; Belle, M.; Domart, M.-C.; Larijani, B.; Collinson, L.; Raposo, G.; Salamero, J. eC-CLEM: Flexible Multidimensional Registration Software for Correlative Microscopies. *Nat Methods* **2017**, *14* (2), 102–103.

(3) Yang, J. E.; Larson, M. R.; Sibert, B. S.; Shrum, S.; Wright, E. R. CorRelator: Interactive Software for Real-Time High Precision Cryo-Correlative Light and Electron Microscopy. *Journal of Structural Biology* **2021**, *213* (2), 107709.

(4) Schmidt, M.; Rohde, F.; Braumann, U.-D. Visualization and Co-Registration of Correlative Microscopy Data with the ImageJ Plug-in Correlia. In *Methods in Cell Biology*; Elsevier, 2021; Vol. 162, pp 353–388.

(5) Stirling, D. R.; Swain-Bowden, M. J.; Lucas, A. M.; Carpenter, A. E.; Cimini, B. A.; Goodman, A. CellProfiler 4: Improvements in Speed, Utility and Usability. *BMC Bioinformatics* **2021**, *22* (1), 433.

(6) Stringer, C.; Wang, T.; Michaelos, M.; Pachitariu, M. Cellpose: A Generalist Algorithm for Cellular Segmentation. *Nat Methods* **2021**, *18* (1), 100–106.

(7) Koning, R. I.; Vader, H.; van Nugteren, M.; Grocutt, P. A.; Yang, W.; Renault, L. L. R.; Koster, A. J.; Kamp, A. C. F.; Schwertner, M. Automated Vitrification of Cryo-EM Samples with Controllable Sample Thickness Using Suction and Real-Time Optical Inspection. *Nat Commun* **2022**, *13* (1), 2985.
